# Sequence-Independent RNA Sensing in Living Mammalian Cells

**DOI:** 10.1101/2025.09.18.677213

**Authors:** Natalie S. Kolber, K. Eerik Kaseniit, Tobias V. Lanz, William H. Robinson, Xiaojing J. Gao

## Abstract

Recently, several groups described sensors in living cells that take advantage of adenosine deaminases acting on RNA (ADARs) to link the presence of an RNA (a “target transcript”) to the translation of a payload from a second, exogenously introduced mRNA. These sensors share the key mechanism of editing a stop codon opposite a specific sequence motif in the target transcript, where this motif requirement is dictated by ADAR’s strong sequence preference. This constrains sensor design and precludes the sensing of short sequences that lack such motifs, often essential for key applications such as sensing viral RNAs and differentiating splice isoforms. Here we address this limitation with modular RNA sensors using adenosine deaminases acting on RNA (“modulADAR”). ModulADAR features two key elements that mirror the modularity of ADARs: regions that hybridize with the target transcript to recruit ADAR’s dsRNA-binding domains, and a stem-loop for stop-codon editing by ADAR’s catalytic domain. We optimize modulADAR and apply it to detect short subsequences that cannot be sensed by prior-generation sensors. We anticipate that modulADAR will empower broader basic science and therapeutic applications, especially those that will uniquely benefit from programmable RNA detection in living cells.

## Main

The abundance of available single-cell transcriptomics data allows a cell’s type and state to be inferred from its RNA signature. However, methods to detect and act upon RNA transcripts in living mammalian cells have historically been lacking. Such tools could enable the identification and manipulation of specific cell types in living organisms for basic research, the selective ablation of pathogenic cells in human patients (*e.g*., cancer cells or cells implicated in autoimmunity), and the cell type-specific delivery of *in vivo* gene therapies. Towards this goal, we alongside other groups recently described a system for RNA sensing leveraging the RNA-editing abilities of adenosine deaminases acting on RNA (ADARs).^1–3^ These sensors take advantage of ADAR’s ability to edit adenosines in double-stranded RNA (dsRNA), altering a stop codon and allowing for the translation of a downstream payload conditioned upon the presence of a specific target transcript that hybridizes with the sensor RNA.

However, the applicability of prior-generation “linear” sensors is fundamentally limited by their dependence on ADARs’ sequence preferences. Linear sensors require two critical elements: a stop codon to prevent constitutive payload expression, and a target-sensor duplex structure that serves as an efficient substrate for ADAR editing (**Supplementary Figure 1**). Specifically, prior work relies on ADARs’ preference for editing adenosines in A:C mismatches within otherwise double-stranded regions, necessitating a CCA motif in the target transcript positioned opposite the sensor’s UAG stop codon (although other sequences have been found to function as suitable motifs albeit with reduced efficiency^3, 4^). This constraint significantly limits sensor optimization: due to the CCA motif requirement, researchers cannot freely select optimal sensing regions with low secondary structure nor empirically test potential target sites through exhaustive tiling. Even more crucially, the CCA requirement can completely prohibit the sensing of short transcripts or subsequences that lack the necessary motif, such as in the case of detecting alternative splicing events that underly many physiologically and pathologically relevant cell states.

Analogous to efforts engineering Cas9 nucleases with broader protospacer-adjacent motif (PAM) specificities^5–7^, we have developed a next-generation strategy that overcomes sequence constraints. We dubbed these new sensors “modular RNA sensors using adenosine deaminases acting on RNA (modulADAR).” ModulADAR is free from target transcript motif constraints, significantly expanding the universe of detectable transcripts. Unlike linear sensors, modulADAR relies on ADAR editing a stem-loop sequence derived from endogenous ADAR substrates. We screen stem-loops to identify those that enable conditional sensor activation, showing that the choice of stem-loop is orthogonal to the target transcript and identifying strategies to optimize signal and baseline. Finally, we demonstrate modulADAR’s potential for sensing viral RNAs and splice isoforms.

## Results

### Foundational engineering of stem-loop sensors

Linear sensors rely on specific trinucleotide motifs in the endogenous target transcript. For example, prior work primarily used a CCA motif to complex with the UAG stop codon and form a substrate preferred by ADARs.^1–3^ In contrast, we posited that mirroring the modularity of ADARs would yield more efficient, sequence unconstrained sensors. Specifically, we designed modulADAR to reflect ADAR’s two functionalities (RNA binding via the dsRNA binding domains, RBDs, and editing by the catalytic deaminase domain, DD).^8^ Sensors comprise an optional, constitutively translated marker protein (*e.g*., mCherry) followed by a region reverse-complementary to the target transcript bisected by a stem-loop. We selected stem-loop sequences derived from endogenous ADAR substrates that are endogenously edited with high efficiency (up to 99%). The stem-loop either already contains or is modified to house a stop codon that prevents the expression of a downstream payload (*e.g*., GFP). As in prior work^1–3^, 2A sequences separate flanking coding sequences from the central sensing region. Here, we expect ADAR to be recruited through its RBDs to the dsRNA region formed between target transcript and sensor, while editing occurs within the stem-loop via the DD (**Figure 1a**). This approach would eliminate the sequence motif requirements that limited previous sensor designs.

**Figure 1.**
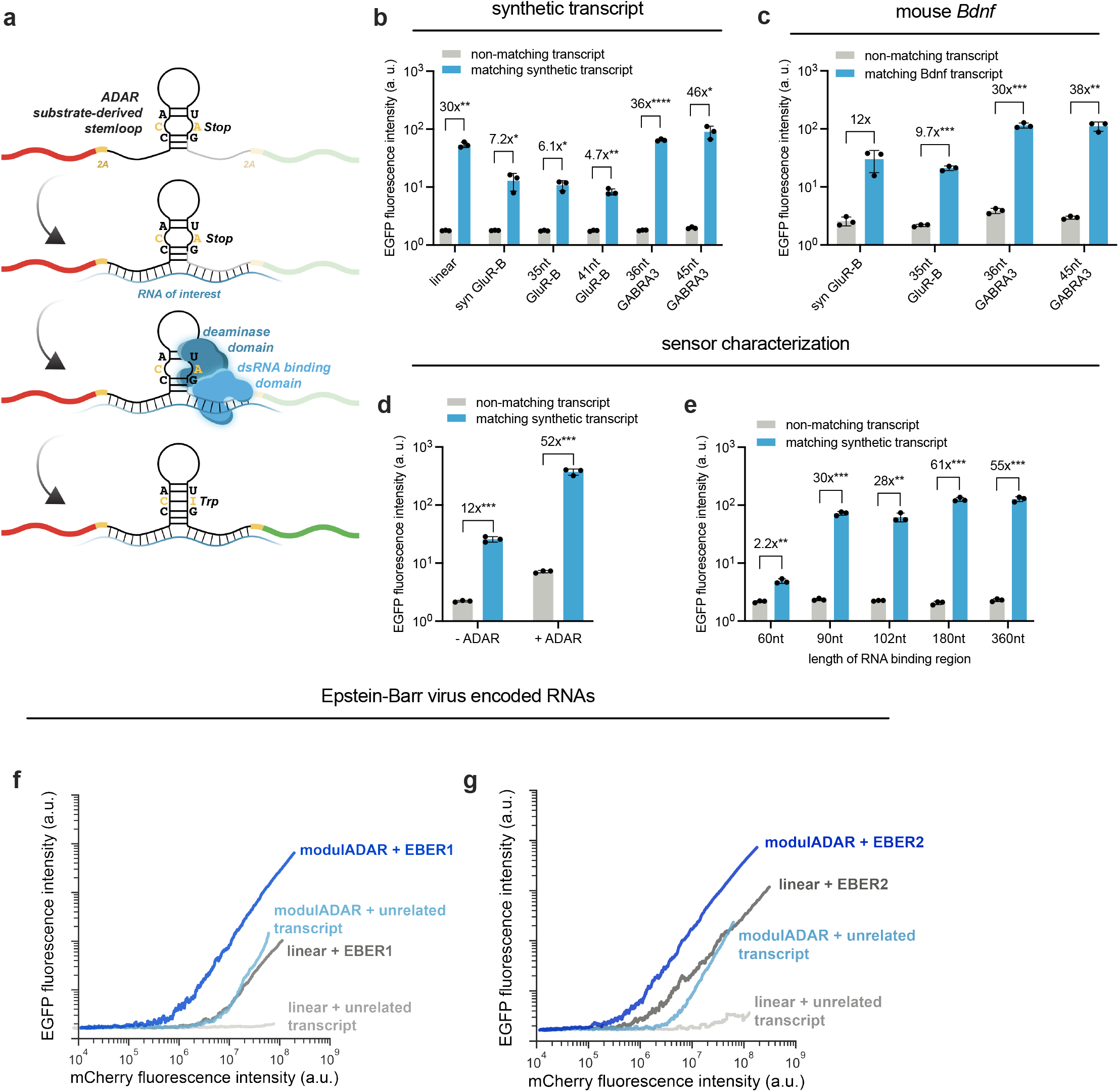
Development of modulADAR towards sensing viral infection. (a) ModulADAR sensors function by separately engaging ADAR’s RNA binding and deaminase domains, using a target-binding region to recruit ADAR to the RNA of interest and a stem-loop structure containing a stop codon for conditional editing and gating translation. (b) Comparison of different stem-loops targeting a synthetic 3’ UTR target sequence. The 45nt GABRA3 stem-loop demonstrates the best signal and fold-activation. (c) Comparison of different stem-loops targeting the 3’ UTR of mouse *Bdnf*. Relative performance of stem-loops is similar to sensors targeting the synthetic target sequence, suggesting orthogonality between the target binding region and the stem-loop.(d) Comparison of modulADAR performance with endogenous ADAR levels versus ADAR1^p150^ overexpression, demonstrating improvement in signal intensity and fold-activation with overexpression. (e) Effect of target RNA binding region length on sensor performance, demonstrating a minimal target binding region length of around 90nt. (f, g) Traces of mean EGFP fluorescence intensity across bins of mCherry (transfection marker) fluorescence intensity. Transfection efficiency increases from left to right while reporter expression increases from bottom to top. ModulADAR sensors generally outperform linear sensors at lower transfection efficiencies.

We investigated modulADAR via transient transfection of sensor and target plasmid DNA (pDNA) in human embryonic kidney (HEK) 293 cells. First, we screened for stem-loops capable of mediating target transcript-dependent activation using a synthetic 3’ UTR target sequence (**Figure 1b)**. We found that GluR-B R/G-site derived sequences as well as GABRA-3 derived sequences^9, 10^ could be used to engineer modulADAR sensors. Sensors using GABRA-3 derived stem-loops perform comparably to linear sensors in this context (**Figure 1b**). We then asked if these stem-loops could be extended towards sensing an unrelated transcript, designing sensors for the 3’ UTR of mouse *Bdnf* (**Figure 1c**). Gratifyingly, the relative signal for the stem-loops tested is similar between these sensors and the previously tested synthetic transcript sensors (**Figure 1c**). This suggests that the choice of stem-loop is orthogonal to the target RNA binding region, such that the target binding region and stem-loop can be optimized independently of one another. Going forward, we therefore used the best-performing, 45nt GABRA3 stem-loop.

Although all human cells naturally express ADAR,^11^ our initial experiments utilized ADAR1^p150^ overexpression based on previous observations that this significantly enhances editing efficiency in linear sensors.^1–3^ Given that ADAR overexpression is suboptimal for therapeutic applications however, we also assessed modulADAR performance with endogenous ADAR levels (**Figure 1d**). While the system functions without exogenous ADAR, we observed that overexpression significantly improves both amplitude and signal-to-baseline ratio (**Figure 1d**). To facilitate more sensitive detection of design improvements, subsequent data were therefore collected with ADAR1^p150^ overexpression. Finally, we determined rules for sensor length, establishing a minimal target RNA binding region length of ~90nt (**Figure 1e**). Stronger activation was achieved with longer sensors of 180-360 bp, consistent with previous reports on linear sensors.^1–3^

### Sensing viral RNAs using modulADAR

One compelling application of RNA sensors is sensing viral infection. This could enable both therapeutic applications (*e.g*., elimination of infected cells via targeted delivery of cytotoxic payloads) as well as basic research through the real-time visualization of RNA expression. The high expression level of many viral transcripts^12^ make them excellent candidates for detection using RNA sensors. However, these RNAs are often short and highly structured, meaning they may lack a conformationally exposed trinucleotide motif suitable for linear sensors.

An example of viral RNAs of clinical interest are the Epstein-Barr virus (EBV)-associated RNAs EBER1 and EBER2. EBV is a common herpesvirus that infects ~95% of people worldwide and establishes lifelong latency in B cells. Latent EBV infection has been linked to multiple sclerosis (MS)^13, 14^ and other autoimmune diseases.^15^ This suggests that eliminating EBV-infected cells could be beneficial. In fact, the efficacy of anti-CD19 CAR-T cell therapy for the treatment of MS^16^ and other autoimmune disorders^17, 18^ may arise due to depletion of EBV^+^ B cells. However, these treatments result in complete B-cell loss, whereas an ideal therapy would target only EBV-infected cells. One possible strategy is to target EBV proteins, but this is limited by the absence of viral protein expression during EBV latency type 0 in resting memory B cells, where only the short noncoding RNAs EBER1 and EBER2 are expressed.^19^

We envisioned that RADAR sensors designed to deploy a cytotoxic payload in the presence of EBER1 or EBER2 could facilitate the selective elimination of infected B cells. As the first step towards this goal, we evaluated modulADAR’s ability to detect EBER1 and EBER2. These RNAs exemplify the need for sequence-unrestricted sensors, as the two sequences contain only a total of three CCA sites. We tiled the EBER1 and EBER2 consensus sequences (GenBank: MG021311.1^20^; 167nt and 173nt in length, respectively) with 90nt-long modulADAR sensors at approximately 10nt intervals. We also designed 90nt-long linear sensors around the two CCA sites in EBER1 and the one CCA site in EBER2. We evaluated these sensors in HEK cells by co-transfecting sensor pDNA alongside either control pDNA or pDNA expressing EBER1/EBER2 transcripts (**Supplementary Figure 2a, b**).

Between the two EBV-associated RNAs, EBER1 represents the more attractive target due to its longer half-life and consequently higher cellular concentration (approximately 10^6^ copies per cell^21^). Our assessment revealed that for EBER1, a modulADAR sensor outperformed linear sensors in both absolute signal and signal-to-baseline ratio (**Supplementary Figure 2a**). Conversely, for EBER2, while a linear sensor showed stronger fold-activation, it generated substantially weaker absolute signal than any modulADAR design (**Supplementary Figure 2b**). In the context of an *in vivo* gene therapy, we expect such low signal might be a liability as delivery efficiency would be substantially lower than in the context of transient transfection. We therefore examined the performance of the best performing EBER1 and EBER2 sensors across a range of transfection efficiencies (**Figure 1f, g**). While linear sensors exhibit better fold-activation in the most highly transfected cells, they display reduced signals in less transfected cells. In contrast, although modulADAR sensors have less fold-activation in the most highly transfected cells, they outperform linear sensors as transfection efficiency decreases (**Figure 1f, g**). The elevated signal of modulADAR is thus more suited than linear RADAR to this potential use case.

Dots represent biological replicates with bars showing group means (n = 3 for a, b, c, d; n = 2 for f, g). Statistical significance was assessed using two-tailed Student’s t-test with Bonferroni correction for multiple comparisons. Significance levels: \*\*\*\**P* < 0.0001, \*\*\**P* < 0.001, \*\**P* < 0.01, \**P* < 0.05 (not shown for f, g).

### Developing sensors with improved fold-activation for distinguishing splice isoforms

A second compelling application of RNA sensing is distinguishing alternative splice isoforms. The vast majority of human genes undergo alternative splicing,^22^ with different isoforms playing unique roles in both normal cellular function and disease pathogenesis. For example, Bcl-x splicing generates either pro- or anti-apoptotic proteins whose balance influences cancer progression,^23^ while neurexins undergo extensive alternative splicing to create thousands of distinct adhesion molecules that specify synaptic connections.^22^ Despite the biological importance of detecting specific splice variants, the relatively short length of exons (median ~120 bp^24^) often precludes the use of linear RADAR sensors, as many exons lack the required trinucleotide motif.

One example of an exon lacking a canonical “CCA” motif required by linear sensors is the 54-bp long exon 7 of human *SMN* (survival of motor neuron). Humans possess two nearly identical copies of the *SMN* gene: *SMN1* and *SMN2*. While both genes contain identical exon 7 sequences, a single nucleotide difference in *SMN2* intron 7 disrupts proper splicing, causing most *SMN2* transcripts to produce a truncated, poorly functional protein (*SMNΔ7*). Spinal muscular atrophy, a rare genetic neuromuscular disorder, is the result of mutations in *SMN1* that cause patients to rely on the only partially functional *SMNΔ7* transcript^25^. Upregulation of alternative splicing to include exon 7 in *SMN2* is therefore the target of the FDA-approved drugs nusinersen^26^ and risdiplam.^27, 28^ A simple live-cell reporter for such splicing events would accelerate the discovery of new splice modulators for many disorders that currently lack effective treatments.

Towards sensing *SMN* exon 7 inclusion, we first evaluated whether alternative trinucleotide motifs (to CCA) in the target transcript could serve as a trinucleotide motif enabling the use of linear sensors. We therefore exhaustively mapped alternative trinucleotides using a 90bp, linear sensor against a synthetic target transcript and identified a general motif (U/G/C)NA that allows for comparable sensing (**Supplementary Figure 3**). As *SMN2* exon 7 contains four such motifs, we designed linear sensors around each site. However, these sensors displayed poor activation and did not outperform the modulADAR sensor (**Figure 2a**). This suggests that alternative trinucleotide motifs exhibit higher context dependency than the canonical CCA, underscoring the value of modulADAR’s sequence-independence.

**Figure 2.**
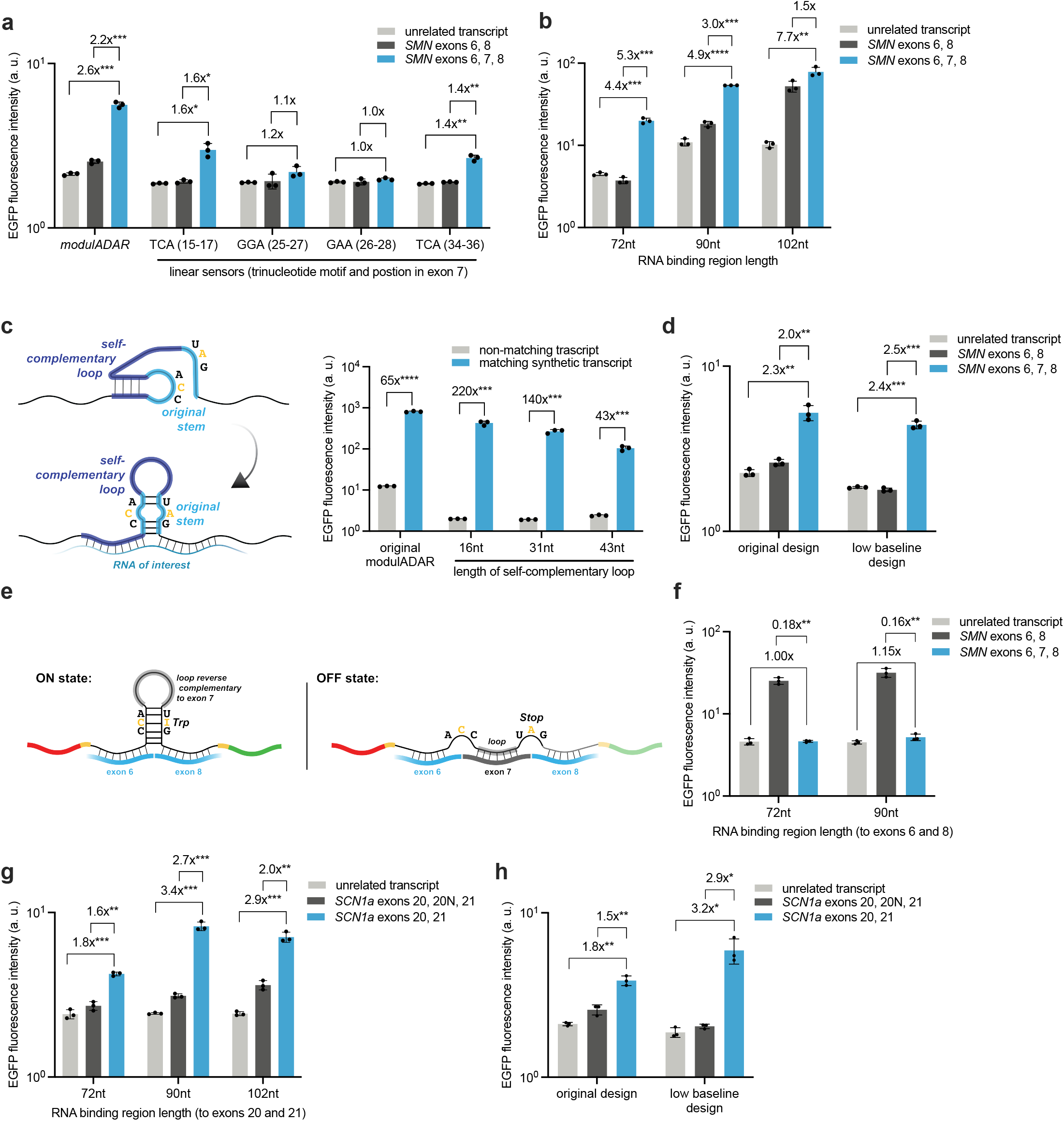
Developing modulADAR sensors for distinguishing spice isoforms. (a) Performance of linear sensors designed around four alternative motifs in *SMN* exon 7, showing poor activation compared to modulADAR. (b) Detection of *SMN* exon 7 inclusion using modulADAR sensors of varying target binding region lengths, with the 72nt sensor showing optimal performance. (c) Self-complementary sensors designed using strand displacement principles enable improved fold-activation while maintaining comparable signal to the original stem-loop design. (d) An optimized self-complementary sensor for *SMN* exon 7 inclusion modestly outperforms the original sensor design. (e) Schematic illustration of an *SMN* exon 7 exclusion sensor design that specifically detects *SMNΔ7* transcripts. (f) Performance of the *SMN* exon 7 exclusion sensor, showing activation in the presence of *SMNΔ7* but not in the presence of *SMN* exon 7 or an unrelated control transcript. (g) Detection of *SCN1a* exon 20N exclusion using various target binding region lengths, with the 90nt sensor demonstrating optimal performance. (h) A self-complementary sensor for *SCN1a* exon 20N exclusion outperforms the original sensor design. Statistical significance was assessed using two-tailed Student’s t-test with Bonferroni correction for multiple comparisons. Significance levels: ****P < 0.0001, ***P < 0.001, **P < 0.01, *P < 0.05.

To tune modulADAR’s ability to detect exon 7 inclusion, we transiently transfected HEK cells with plasmids encoding modulADAR sensors of varying target binding region lengths along with plasmids expressing either an unrelated transcript, *SMN* exons 6 and 8 (Δ7), or *SMN* exons 6, 7, and 8 (full-length) (**Figure 2b**). The shortest sensor tested (72nt) performed best, likely because this sensor has the least overlap with exons 6 and 8 (**Figure 2b**).

For applications in small molecule discovery, we envision that human cells cultured *in vitro* would be transduced or transiently transfected with the relevant sensor prior to compound dosing. The copy number of sensor would therefore be significantly higher than the copy number relevant to *in vivo* gene therapies. At the same time, high absolute signal as well as high fold-activation would better facilitate compound screening. This is a distinct scenario from the signal-limited viral detection mentioned above. We therefore sought strategies to mitigate baseline signal while maintaining high absolute signal.

We hypothesized that baseline signal might result from the ability of the short double-stranded region in the stem-loop to recruit ADAR independently of target transcript. We therefore explored strategies to reduce this non-specific activity. First, we designed sensors with an additional stop codon, either by editing the stem-loop to contain an extra stop codon or by extending the stem-loop by an additional stop codon (**Supplementary Figure 4**). While these sensors exhibit lower baseline, signal is also depressed, resulting in activation that is inferior to the original sensor design (**Supplementary Figure 4**).

Our second approach was based on principles of strand displacement.^29^ We engineered sensors with a self-complementary region that replaces the conventional loop and hybridizes with the sensor’s target binding region (**Figure 2c**). In the absence of the target transcript, we expect the sensor to adopt an alternative folding conformation where the self-complementary region binds to the target binding region, reducing the formation of the stem-loop structure required for ADAR editing. However, when the target transcript is present, it competitively binds to the target binding region, displacing the self-complementary segment. This enables the formation of the functional stem-loop structure required for ADAR editing while simultaneously providing the dsRNA region for RBD recruitment.

We evaluated sensors with self-complementary loops of different lengths. The longest version tested demonstrated reduced signal intensity and inferior performance compared to the original sensor design. In contrast, shorter variants maintained signal intensity comparable to the original while exhibiting reduced baseline activation, yielding improved fold-activation (**Figure 2c**). We interrogated even shorter loop lengths, observing that shorter loops generally led to higher baseline, consistent with our proposed strand displacement mechanism (**Supplementary Figure 5a**). Notably, this offers a strategy for rationally tuning both absolute and baseline signal. Finally, we tested different configurations of the self-complementary sequences within the loop structure in relation to the target binding region but found that each performed similarly to the original self-complementary design (**Supplementary Figure 5b, c**). We then applied this principle towards designing better *SMN* exon 7 sensors. We screened self-complementary, low baseline versions of the modulADAR *SMN* sensor (**Supplementary Figure 6**), identifying a sensor that modestly outperforms the original design (**Figure 2d**).

For some applications, we are more interested in the absence of a particular exon rather than its presence. We therefore created a sensor that activates specifically in response to *SMNΔ7*, inspired by the self-complementary loop design described above. Here, the target binding region is reverse complementary to exons 6 and 8, while the original loop is replaced with a region reverse complementary to exon 7. In the presence of the full-length isoform, the loop binds to exon 7, preventing the formation of the stem-loop critical for ADAR editing (**Figure 2e**). This design enables sensors that activate in the presence of *SMNΔ7*, but not in the presence of an unrelated transcript or the isoform containing exon 7 (**Figure 2f**).

The *SCN1a* gene presents a more clinically relevant example for such activation by exon exclusion. Haploinsufficiency in *SCN1a* causes Dravet syndrome, the most common form of genetic epilepsy, characterized by frequent, severe seizures, developmental delays, and an increased risk of sudden unexpected death in epilepsy (SUDEP).^30^ The *SCN1a* gene occasionally incorporates a “toxic” 20N exon during splicing, which introduces a premature stop codon that destabilizes the transcript through nonsense-mediated decay. Strategies for treating Dravet therefore include an anti-sense oligonucleotide (ASO) that prevents exon 20N inclusion, thereby increasing the expression of functional *SCN1a*.^31^

To detect exon 20N exclusion, we designed sensors with a loop reverse complementary to the entire 20N exon (64nt). We validated our sensor designs by co-transfecting HEK cells with sensors of varying target binding region lengths alongside either an unrelated control transcript, *SCN1a* exons 20, 20N, and 21, or *SCN1a* exons 20 and 21 (**Figure 2g**). We found that the sensor with a 90nt target binding region performed the best (**Figure 2g**). Building on our low baseline sensor design, we then incorporated self-complementary sequences at the 5’ end of the loop that could hybridize with the sensor’s target binding region (**Supplementary Figure 7a**). After screening sensors with different self-complementary region lengths (**Supplementary Figures 7b, c**), we identified a sensor that outperforms the original design (**Figure 2h**).

### Discussion

In this work, we improve on previous ADAR-dependent RNA sensors with modulADAR, mirroring the functional and structural modularity of ADARs to enable sequence-unconstrained RNA sensing. Unlike prior-generation sensors, this approach lacks fundamental sequence constraints, thereby expanding the range of detectable transcripts. We identify stem-loops most amenable to sensor design and characterize design features such as target binding region length and strategies for minimizing target-independent activation. We apply modulADAR to sensing transcripts that are not readily amenable to prior-generation designs, including viral RNAs and splice isoforms.

While modulADAR overcomes key limitations of prior sensors, important challenges remain to be addressed in future work. Firstly, all experiments described were performed by transient transfection of sensor and target transcripts, and we have not yet demonstrated sensing of endogenous RNAs. This may require further optimization, as endogenous RNAs are typically expressed at lower levels than those achieved through transient transfection and we believe that the probability of productive sensor-transcript interactions is concentration-dependent and diffusion-limited. To mitigate this, we are developing strategies to increase sensor and target RNA co-localization beyond sequence complementarity.

Further stem-loop engineering has the potential to address both sensor performance without ADAR overexpression and improve the detection of low-abundance transcripts. While our initial experiments used stem loops derived from natural substrates, rational design enabled sensors with better fold-activation, suggesting that evolution has not fully optimized these elements for our application. High-throughput screens therefore have the potential to further optimize stem-loop design.

Importantly, ideal stem-loop characteristics will vary by use-case. For example, *in vivo* gene therapies for killing cancerous or virally-infected cells require low target-independent activation but may not require high payload expression, especially if using highly potent effectors like AB toxins.^32^ Similarly, as Cre is used routinely for research applications,^33^ it was previously demonstrated that Cre significantly increases the signal from linear sensors.^1–3^ These applications could be readily enabled by our strategy for rationally reducing baseline using self-complementary sensors. Conversely, applications such as *in vivo* CAR-T therapy require robust output expression in target cells while potentially tolerating modest off-target activation. Our finding that stem-loop performance remains consistent across different target transcripts suggest that principles for tuning both baseline and signal established through high-throughput screens will be broadly applicable and enable rapid optimization for diverse applications.

Our demonstration that modulADAR can detect specific splice isoforms suggests therapeutic applications beyond genetic medicines. Traditionally, splicing has been therapeutically modulated using ASOs. However, recently it has been demonstrated that small molecules (which exhibit oral bioavailability and easier manufacturing) can also selectively modulate splicing. This has been exemplified by the approval of risdiplam for treating spinal muscular atrophy. Small molecules that modulate splicing are now being investigated for a variety of indications, including Alzheimer’s,^34^, Huntington’s disease,^35^ and cancers.^36^ While these compounds promise to significantly expand the druggable proteome, current strategies for screening small molecules that influence splicing are lacking. *In vitro* binding assays are poorly predictive of functional changes to RNA splicing^37^ and phenotypic assays typically rely on artificial reporter systems that may not reflect native splicing regulation.^38^

We envision that modulADAR could address this by enabling high-throughput screening of small molecule splicing modulators in physiologically relevant contexts. Encouragingly, this application bypasses many current limitations of our system, as sensors could be expressed at high levels through transient transfection or stable integration. Potential safety risks associated with ADAR overexpression in the context of gene therapies are not applicable, and cytoplasmic ADAR variants could be used to prevent interference with nuclear splicing machinery. Finally, sensitive readouts like luciferase could provide high dynamic range for effective screening. Overall, we anticipate that modulADAR will enable a wide variety of applications inaccessible by prior-generation designs, including in basic science, next-generation genetic medicines, and discovery of small molecule therapeutics.

## Methods

### Plasmid design and construction

All plasmids were constructed using standard restriction-ligation cloning (Thermo Scientific FastDigest Buffer, cat. no. B64, Thermo Scientific FastDigest Enzymes, Thermo Scientific Rapid DNA Ligation Kit, cat. no. K1423).

Sensor plasmids are driven by the SFFV promoter as described in previous work.^2^ Sensor insert sequences were generated by annealing and phosphorylation (NEB T4 ligation buffer, cat no. B0202S, Thermo Scientific T4 polynucleotide kinase, cat. no. EK0032) of synthetic oligonucleotides (IDT). Sensors often differ from perfect reverse-complementarity to the target RNA by a few point mutations introduced to avoid stop codons. Generally, a small number of point mutations does not appear to affect sensor performance.

Target plasmids are generally driven by the CMV promoter. EBER1 and 2 target sequences (GenBank: MG021311.1^20^, 6632-6798 and 6959-7131) were expressed under the control of a U6 promoter, with inserts generated by annealing and phosphorylating synthetic oligonucleotides (IDT). *SMN* and *SCN1a* sequences were commercially synthesized (Twist). The *SMN* target plasmids were based on a previously established reporter system, with a single “A” insertion at position 49 of exon 7 differentiating it from the endogenous sequence.^27^ *SCN1a* sequences were obtained from published work.^39^

### Tissue culture

Cell culture was performed using wild-type HEK293 cells (ATCC CRL-1573) regularly tested for mycoplasma contamination. Cells were maintained at 37°C in a humidified atmosphere containing 5% CO2. Growth medium consisted of Dulbecco’s Modified Eagle Medium (DMEM, Fisher Scientific cat. no. 501015428) supplemented with 10% fetal bovine serum (Thermo Fisher Scientific cat. no. FB12999102), 1 mM sodium pyruvate (EMD Millipore cat. no. TMS-005-C), 2 mM L-glutamine (Genesee Scientific cat. no. 25-509), 1x penicillin-streptomycin (Genesee Scientific cat. no. 25-512), and 1x MEM non-essential amino acids (Genesee Scientific cat. no. 25-536).

### Transient transfection and flow cytometry

HEK293 cells were seeded in 24-well plates and maintained until reaching 70-90% confluency. Cells were transfected with jetOPTIMUS DNA transfection reagent (Polyplus cat. no. 117-15) as per manufacturer’s instructions. Per well, 50 ul of jetOPTIMUS buffer, 0.375 µl of JetOptimus reagent, and 500 ng of pDNA was used. A typical experiment used 200 ng of sensor pDNA, 270 ng of target pDNA, and 30 ng of CAG-ADAR1^p150^ pDNA.

48 hours post-transfection, cells were harvested by trypsinization and resuspended in flow buffer (Hank’s Balanced Salt Solution (HBSS), 1X, without calcium, magnesium, phenol red, cat. no. 95053-196, with 2.5 mg/mL bovine serum albumin). Cells were strained through a 40um nylon mesh (VWR cat. no. 75799-940). Flow cytometry was performed using a Biorad ZE5 Cell Analyzer. Data were analyzed using the cytoflow Python package. Cells were gated for cells, singlets, and highly transfected cells (99.5th to 99.9th percentile mCherry of the lowest-transfected well within an experiment; **Supplementary Figure 8**). This gating strategy yields higher baseline activation and lower fold-activation than would be observed when analyzing a wider band of transfected cells (e.g., all mCherry-positive cells). Restricting analysis to highly transfected cells was chosen because it provides a more stringent test of sensor performance, enabling more effective engineering optimization. Due to variations in transfection efficiency, different experiments may be gated slightly differently, making intra-experiment comparisons more reliable than inter-experiment comparisons.

## Acknowledgements

This work was funded by the National Institutes of Health (NIH; R21EB033858 and DP2EB035891 to XJG, R01AI173189 to TVL and WHR), Longevity Impetus grants (to XJG), Stanford Cancer Institute innovative award (to XJG), Stanford Bio-X Interdisciplinary Initiatives seed grant program (R11-6, to XJG), Wu Tsai Neuroscience Institute seed grant (to XJG), National Science Foundation GRFP (to NSK), Stanford ChEM-H CBI training program (to NSK), EDGE Doctoral Fellowship Program (to NSK), and Stanford Bio-X fellowship (to KEK). We thank J.B. Li for plasmids encoding human ADAR. We thank all Gao lab members for their advice and feedback.

## Data availability

A comprehensive list of plasmids used alongside links to plasmid maps are available in the supplementary information. Select plasmids are deposited in Addgene. Additional plasmids are available upon request.

Reported data are available upon request.

## Contributions

NSK, KEK, TVL, XJG, and WHR designed the study. NSK performed and analyzed most of the experiments, with support from KEK. NSK wrote the manuscript with input from all authors.

## Supplementary information

**Supplementary Figure 1.**
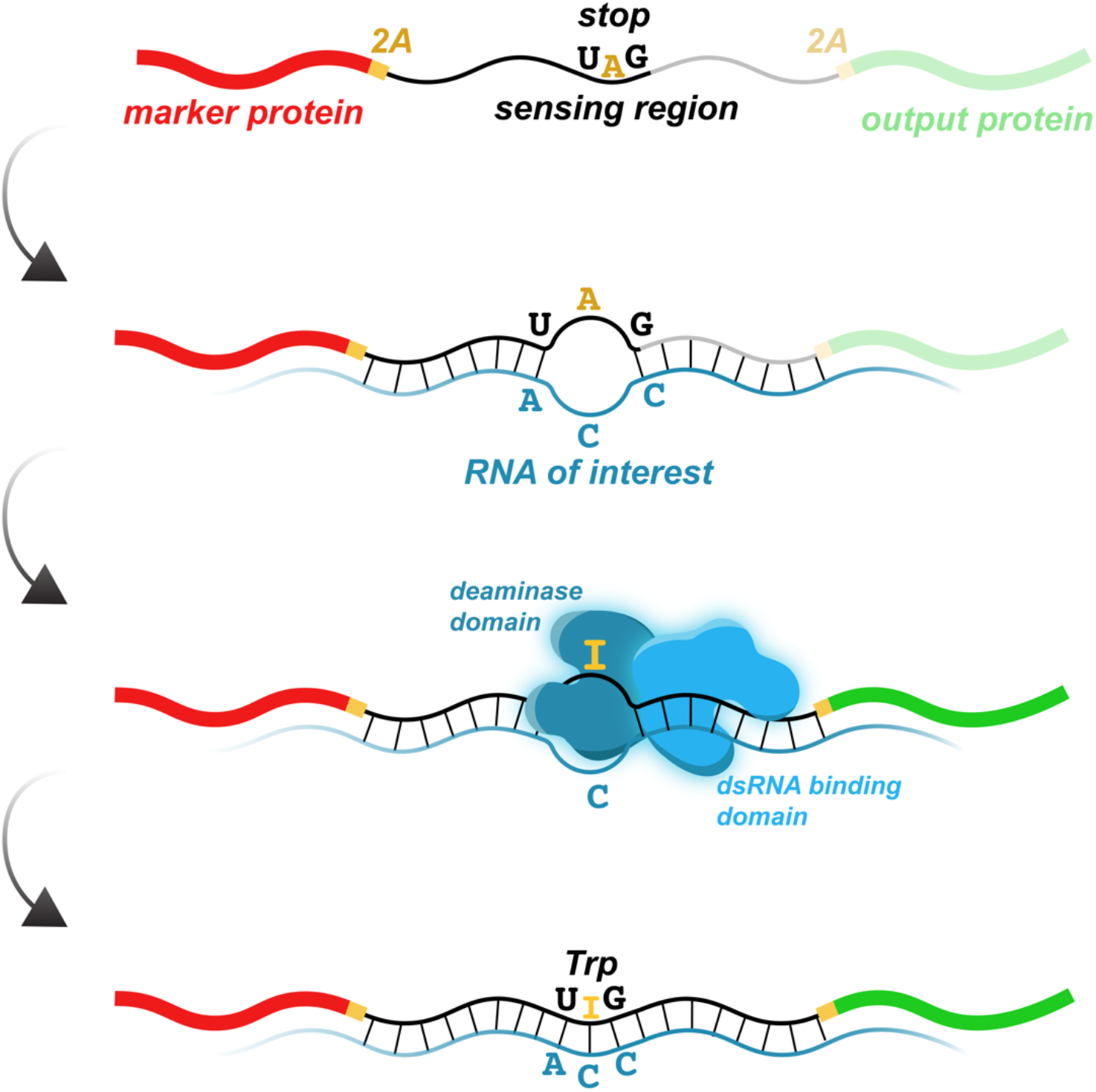
Diagram of prior-generation, linear sensors. Sensors comprise a sensing region reverse-complementary to the target transcript except for a A:C mismatch at a central stop codon, which prevents the translation of the downstream payload. In the presence of the target transcript, dsRNA is formed that binds ADAR by its dsRNA binding domain. ADAR’s deaminase domain deaminates the adenosine in the stop codon to inosine, which is read as guanosine to form a tryptophan codon and enable translation of the payload.

**Supplementary Figure 2.**
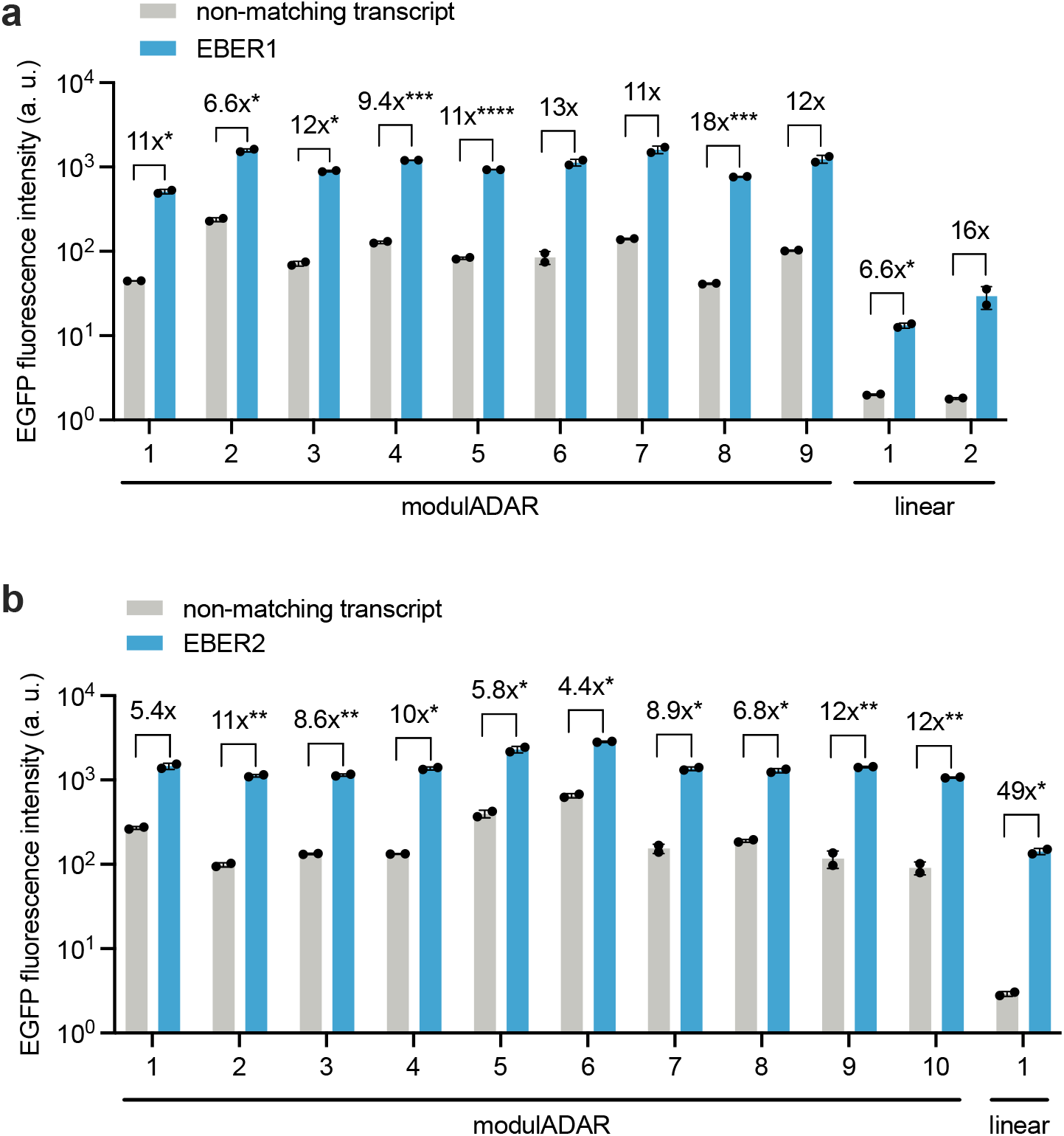
Screening sensors for EBV-encoded RNAs. (a) Detection of Epstein-Barr virus (EBV) EBER1 RNA using modulADAR and linear sensors. Tiling of the EBER1 sequence with modulADAR sensors reveals improved signal and fold-activation compared to linear sensors designed around CCA motifs. (b) Tiling of the EBER2 sequence reveals a linear sensor with improved fold-activation but worse signal than modulADAR sensors. Dots represent biological replicates with bars showing group means (n = 2). Statistical significance was assessed using two-tailed Student’s t-test with Bonferroni correction for multiple comparisons. Significance levels: \*\*\*\**P* < 0.0001, \*\*\**P* < 0.001, \*\**P* < 0.01, \**P* < 0.05.

**Supplementary Figure 3.**
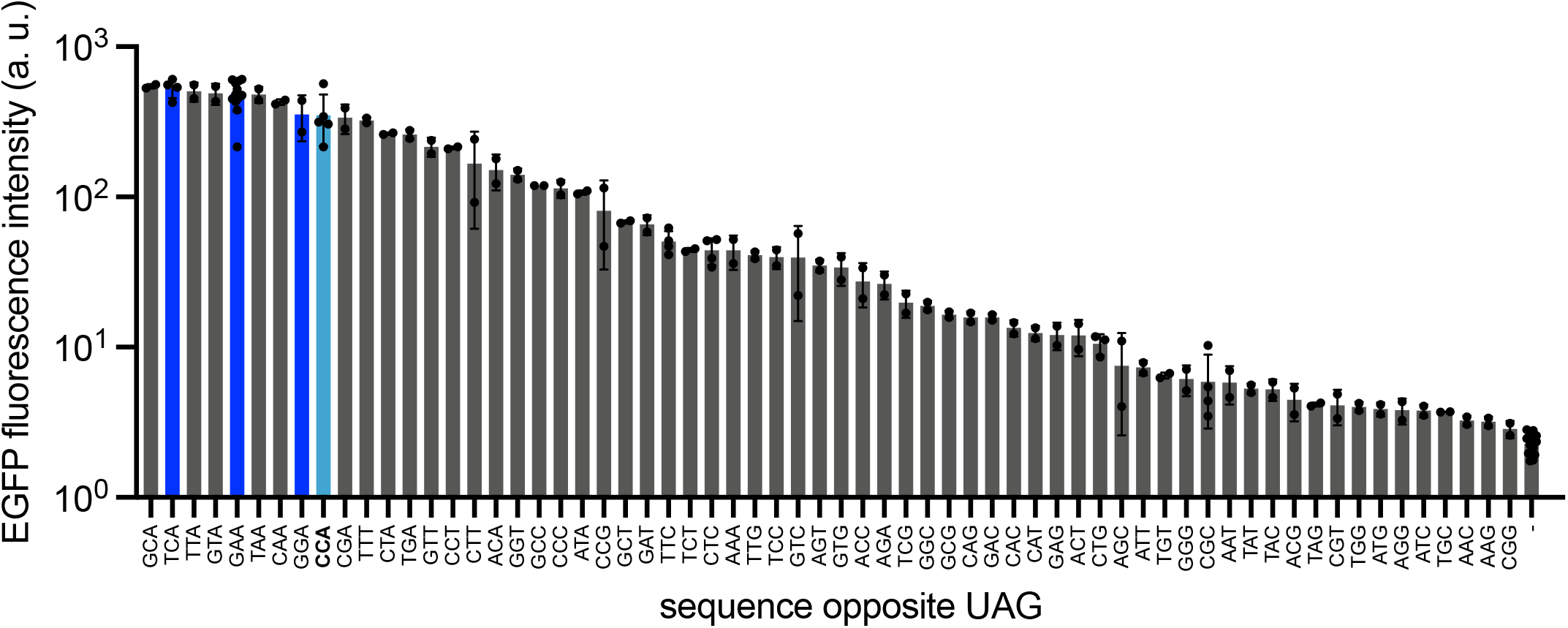
Evaluating alternative trinucleotide motifs for linear sensors. (a) Exhaustive mapping of alternative trinucleotides using a 90nt linear sensor against a synthetic 3’ UTR target transcript. Data suggest a general motif U/G/C)NA that enables comparable sensing to CCA. The original “CCA” motif is highlighted in light blue, and alternative motifs used for *SMN* exon 7 detection are highlighted in dark blue. “-” = non-matching transcript. n = 2-14.

**Supplementary Figure 4.**
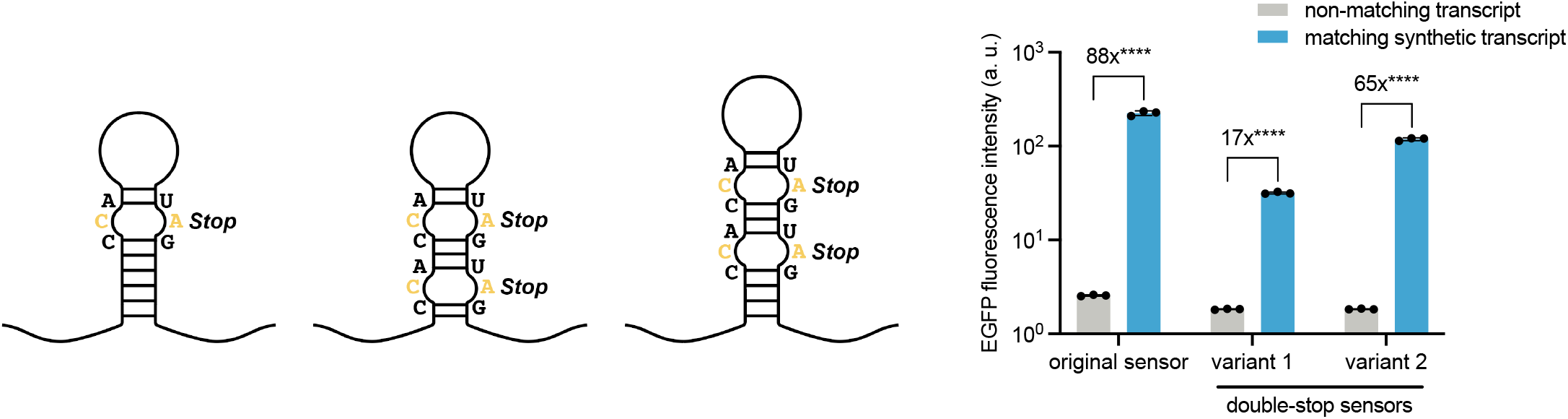
Mitigating baseline signal using multiple stop codons. Stem-loops are designed with an additional stop codon, either by editing the stem-loop to contain an extra stop codon (“double-stop variant 1”) or by extending the stem-loop by an additional stop codon (“double-stop variant 2”). Double-stop stem-loop variants exhibit lower baseline but also reduced fold-activation. Dots represent biological replicates with bars showing group means (n = 3). Statistical significance was assessed using two-tailed Student’s t-test with Bonferroni correction for multiple comparisons. Significance levels: \*\*\*\**P* < 0.0001, \*\*\**P* < 0.001, \*\**P* < 0.01, \**P* < 0.05.

**Supplementary Figure 5.**
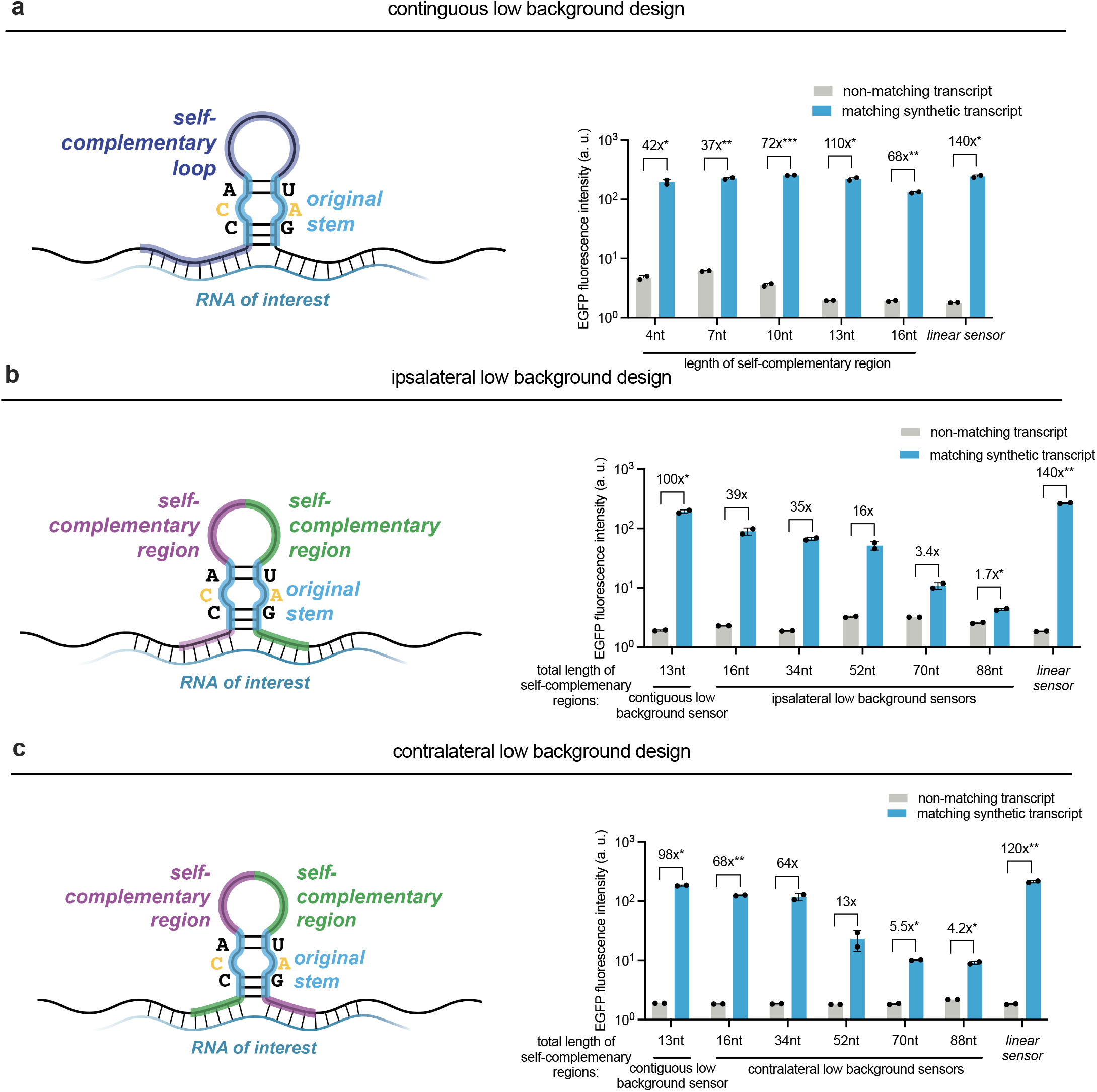
Optimization of self-complementary loop designs. (a) Effect of self-complementary loop length on sensor baseline and activation, showing that shorter loops generally lead to higher baseline activation while longer loops generally lead to decreased signal. (b), (c) Alternative configurations of the self-complementary sequences within the loop, demonstrating similar performance. Dots represent biological replicates with bars showing group means (n = 2). Statistical significance was assessed using two-tailed Student’s t-test with Bonferroni correction for multiple comparisons. Significance levels: ****P < 0.0001, ***P < 0.001, **P < 0.01, *P < 0.05.

**Supplementary Figure 6.**
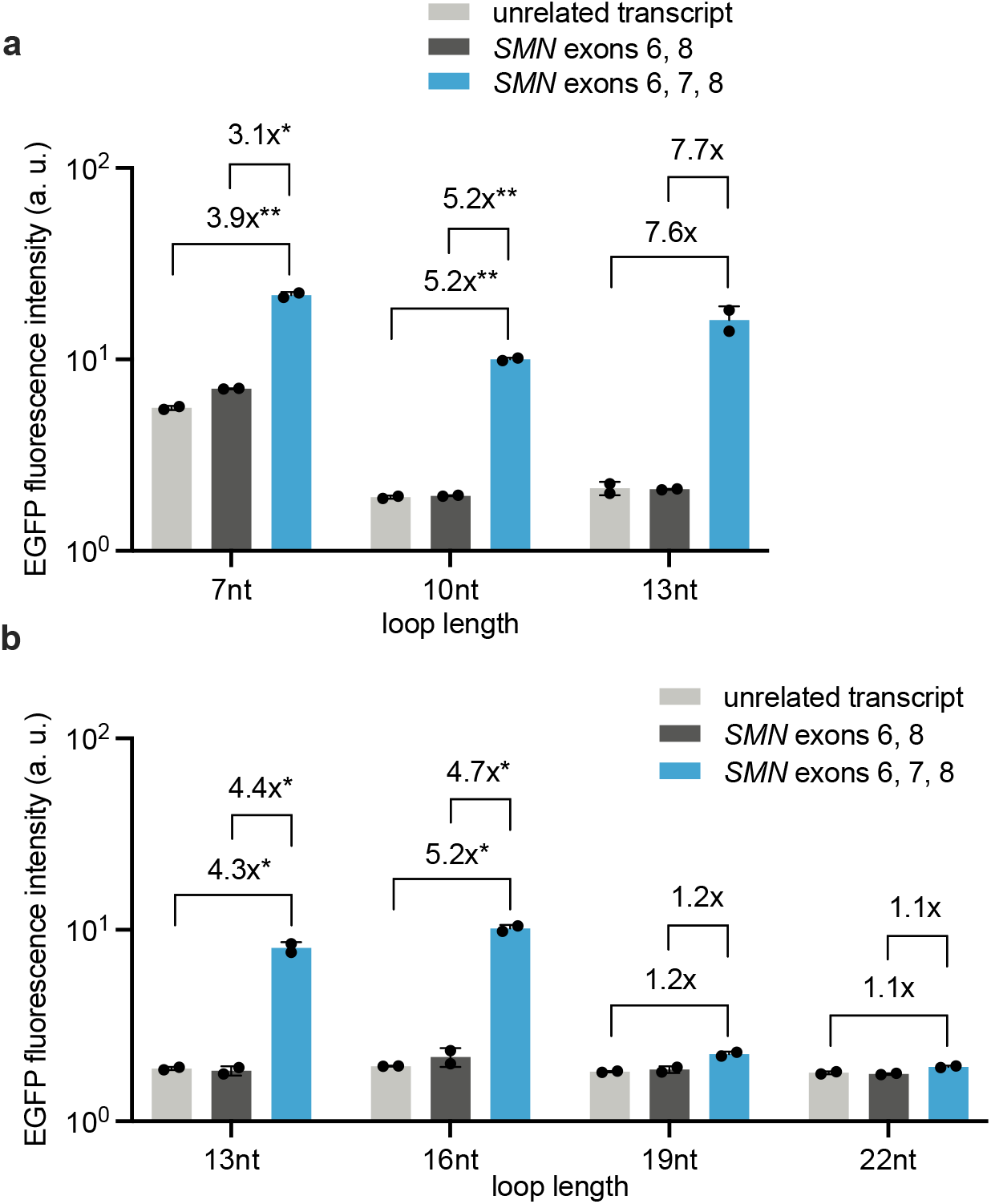
Screening self-complementary, low baseline modulADAR sensors for *SMN* exon 7 detection. (a), (b) Screening of different loop-lengths suggests the optimal sensor has a 16nt-long self-complementary loop. Dots represent biological replicates with bars showing group means (n = 3). Statistical significance was assessed using two-tailed Student’s t-test with Bonferroni correction for multiple comparisons. Significance levels: ****P < 0.0001, ***P < 0.001, **P < 0.01, *P < 0.05.

**Supplementary Figure 7.**
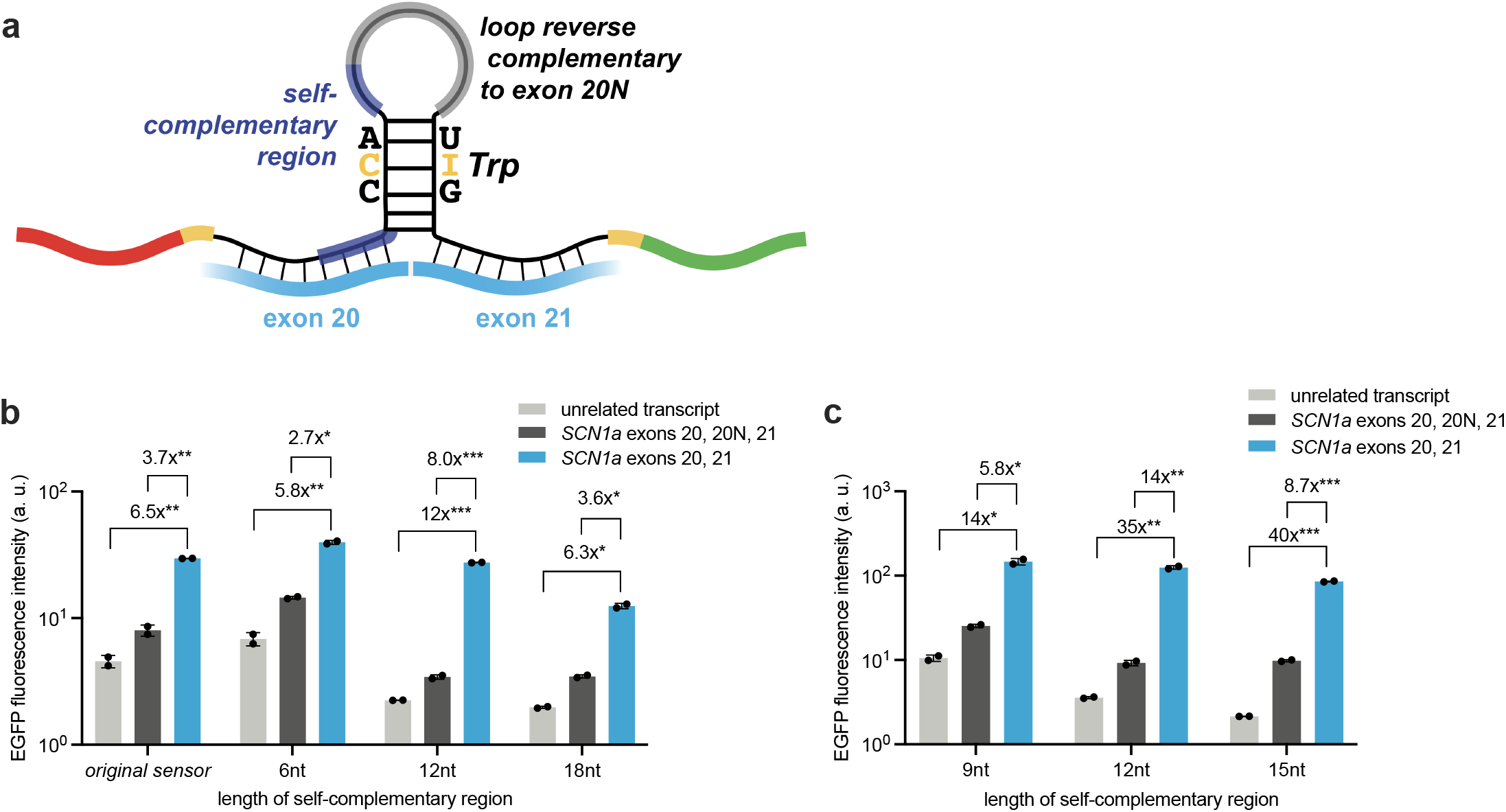
Screening self-complementary, low baseline modulADAR sensors for *SCN1a* exon 20N exclusion. (a) Schematic for low-baseline, self-complementary exon exclusion sensors. A self-complementary sequence is appended at the 5’ end of the loop. (b), (c) Screening sensors with different self-complementary region lengths reveals an optimal sensor with a 12nt self-complementary region. Dots represent biological replicates with bars showing group means (n = 3). Statistical significance was assessed using two-tailed Student’s t-test with Bonferroni correction for multiple comparisons. Significance levels: ****P < 0.0001, ***P < 0.001, **P < 0.01, *P < 0.05.

**Supplementary Figure 8.**
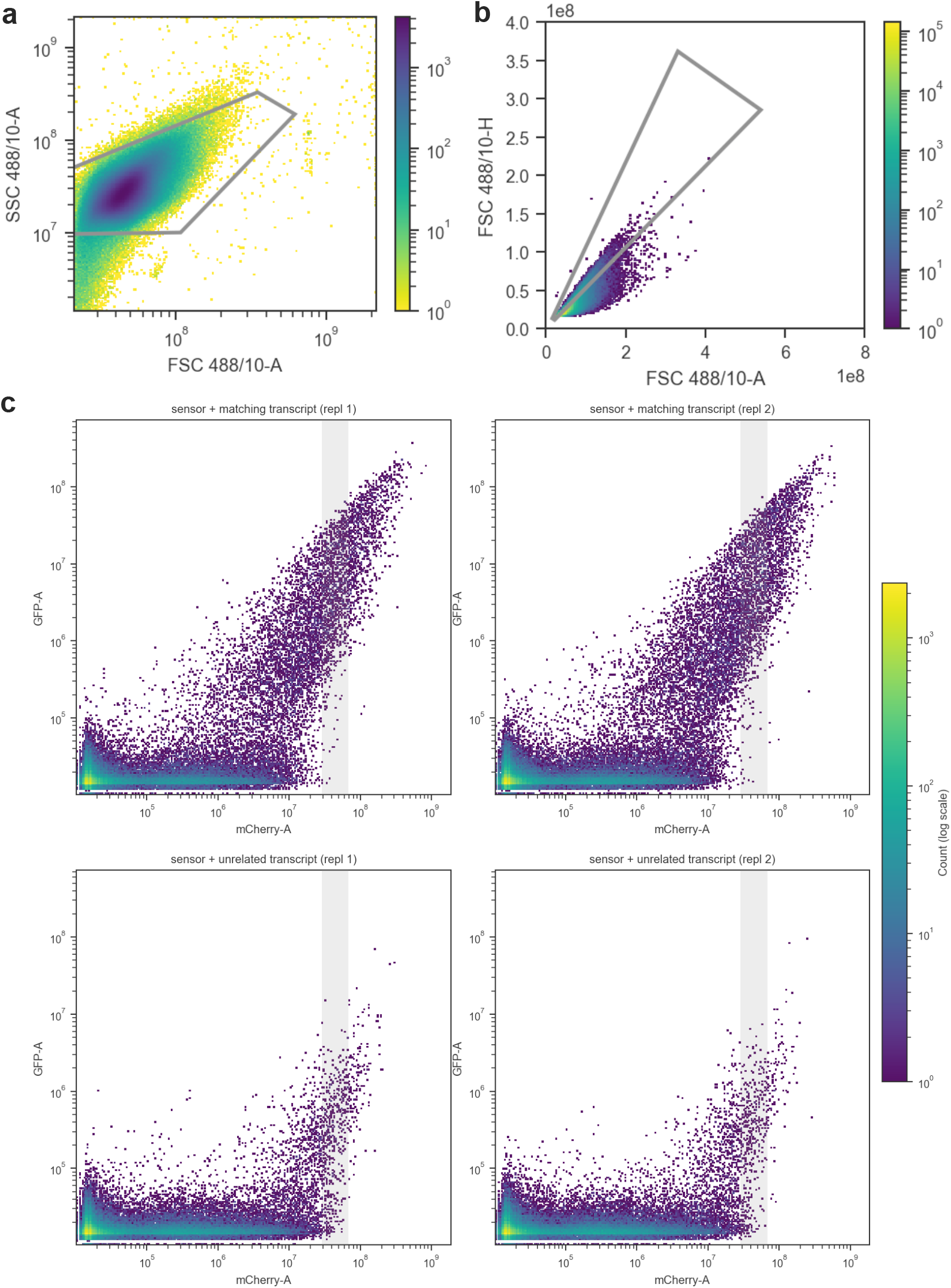
Gating strategy. (a) gating on cells. (b) gating on singlets. (c) Gating on a narrow band of highly transfected cells, indicated by gray bar (99.5th to 99.9th percentile mCherry of the lowest-transfected well within an experiment).

**Supplementary Table 1.** Details of transfections for main text figures.

| Figure | Description | Plasmid | Amount (ng) |
| --- | --- | --- | --- |
| 1b | non-matching transcript | filler plasmid | 270 |
|  | matching synthetic transcript | EK0720 CMV-TO-BFPmut-3UTR(t70587-long) (FLP-IN) | 270 |
|  | linear | EK0438 SFFV-mCherry-SGc(s70587)cSG-EGFP-(3xMS2) (FLP-IN) | 200 |
|  | syn GluR-B | EK0656 SFFV-mCherry-SGc(70587.os4)cSG-EGFP-(3xMS2) | 200 |
|  | 35nt GluR-B | EK0716 SFFV-mCherry-SGc(s70587.os18)cSG-EGFP-(3xMS2) (FLP-IN) | 200 |
|  | 41nt GluR-B | EK0717 SFFV-mCherry-SGc(s70587.os19)cSG-EGFP-(3xMS2) (FLP-IN) | 200 |
|  | 36nt GABRA3 | EK0718 SFFV-mCherry-SGc(s70587.os20)cSG-EGFP-(3xMS2) (FLP-IN) | 200 |
|  | 45ntGABRA3 | EK0698 SFFV-mCherry-SGc(s70587.os13)cSG-EGFP-(3xMS2) (FLP-IN) | 200 |
|  | ADAR overexpression (all conditions) | EK0574 CAG-ADAR1-p150 | 30 |
| 1c | non-matching transcript | filler plasmid | 270 |
|  | matching Bdnf transcript | EK0307 CMV-TO-BFPmut-stop-3UTR(Bdnf-v3) (FLP-IN) | 270 |
|  | linear | EK0301 SFFV-Che-SG(Bdnf.s1)SG-GFP (FLP-IN) | 200 |
|  | syn GluR-B | EK0706 SFFV-mCherry-SGc(Bdnf.os4)cSG-EGFP-(3xMS2) (FLP-IN) | 200 |
|  | 35nt GluR-B | NSK166 SFFV-mCherry-SGc(Bdnf.os18)cSG-EGFP-(3xMS2) (FLP-IN) | 200 |
|  | 36nt GABRA3 | NSK167 SFFV-mCherry-SGc(Bdnf.os20)cSG-EGFP-(3xMS2) (FLP-IN) | 200 |
|  | 45nt GABRA3 | NSK165 SFFV-mCherry-SGc(Bdnf.os13)cSG-EGFP-(3xMS2) (FLP-IN) | 200 |
|  | ADAR overexpression (all conditions) | EK0574 CAG-ADAR1-p150 | 30 |
| 1d | non-matching transcript | filler plasmid | 270 |
|  | matching synthetic transcript | EK0720 CMV-TO-BFPmut-3UTR(t70587-long) (FLP-IN) | 270 |
|  | sensor | EK0698 SFFV-mCherry-SGc(s70587.os13)cSG-EGFP-(3xMS2) (FLP-IN) | 200 |
|  | + ADAR | EK0574 CAG-ADAR1-p150 | 30 |
| 1e | non-matching transcript | filler plasmid | 270 |
|  | matching synthetic transcript | EK0720 CMV-TO-BFPmut-3UTR(t70587-long) (FLP-IN) | 270 |
|  | 60nt | NSK160 SFFV-mCherry-SGc(s70587.os-60bp)cSG-EGFP-(3xMS2) (FLP-IN) | 200 |
|  | 90nt | EK0698 SFFV-mCherry-SGc(s70587.os13)cSG-EGFP-(3xMS2) (FLP-IN) | 200 |
|  | 102nt | NSK159 SFFV-mCherry-SGc(s70587.os-102bp)cSG-EGFP-(3xMS2) (FLP-IN) | 200 |
|  | 180nt | NSK158 SFFV-mCherry-SGc(s70587.os-180bp)cSG-EGFP-(3xMS2) (FLP-IN) | 200 |
|  | 360nt | NSK157 SFFV-mCherry-SGc(s70587.os-360bp)cSG-EGFP-(3xMS2) (FLP-IN) | 200 |
|  | ADAR overexpression (all conditions) | EK0574 CAG-ADAR1-p150 | 30 |
| 1f | unrelated transcript | filler plasmid | 270 |
|  | EBER1 | NSK176 U6TO EBER1 | 270 |
|  | modulADAR | NSK185 SFFV-mCherry-SGc(EBER1.os13.8)cSG-EGFP-(3xMS2) (FLP-IN) | 200 |
|  | linear | NSK266 SFFV-mCherry-SGc(EBER1.linear.2)cSG-EGFP-(3xMS2) (FLP-IN) | 200 |
|  | ADAR overexpression (all conditions) | EK0574 CAG-ADAR1-p150 | 30 |
| 1g | unrelated transcript | filler plasmid | 270 |
|  | EBER2 | NSK177 U6TO EBER2 | 270 |
|  | modulADAR | NSK195 SFFV-mCherry-SGc(EBER2.os13.9)cSG-EGFP-(3xMS2) (FLP-IN) | 200 |
|  | linear | NSK267 SFFV-mCherry-SGc(EBER2.linear.1)cSG-EGFP-(3xMS2) (FLP-IN) | 200 |
|  | ADAR overexpression (all conditions) | EK0574 CAG-ADAR1-p150 | 30 |
| 2a | unrelated transcript | filler plasmid | 270 |
|  | SMN exons 6, 8 | NSK162 CMV-TO-SMN2 minigene Nluc - inactive (exon 6, 8) | 270 |
|  | SMN exons 6, 7, 8 | NSK163 CMV-TO-SMN2 minigene Nluc - active (exon 6, 7, 8) | 270 |
|  | TCA (15-17) | NSK204 SFFV-mCherry-SGc(SNM2.scram1)cSG-EGFP-(3xMS2) (FLP-IN) | 200 |
|  | GGA (25-27) | NSK205 SFFV-mCherry-SGc(SNM2.scram2)cSG-EGFP-(3xMS2) (FLP-IN) | 200 |
|  | GAA (26-28) | NSK206 SFFV-mCherry-SGc(SNM2.scram3)cSG-EGFP-(3xMS2) (FLP-IN) | 200 |
|  | TCA (34-36) | NSK207 SFFV-mCherry-SGc(SNM2.scram4)cSG-EGFP-(3xMS2) (FLP-IN) | 200 |
|  | ADAR overexpression (all conditions) | EK0574 CAG-ADAR1-p150 | 30 |
| 2b | unrelated transcript | filler plasmid | 270 |
|  | SMN exons 6, 8 | NSK162 CMV-TO-SMN2 minigene Nluc - inactive (exon 6, 8) | 270 |
|  | SMN exons 6, 7, 8 | NSK163 CMV-TO-SMN2 minigene Nluc - active (exon 6, 7, 8) | 270 |
|  | 72nt | NSK173 SFFV-mCherry-SGc(Smn2ex7.os13-72bp)cSG-EGFP-(3xMS2) (FLP-IN) | 200 |
|  | 90nt | NSK171 SFFV-mCherry-SGc(Smn2ex7.os13-90bp)cSG-EGFP-(3xMS2) (FLP-IN) | 200 |
|  | 102nt | NSK172 SFFV-mCherry-SGc(Smn2ex7.os13-102p)cSG-EGFP-(3xMS2) (FLP-IN) | 200 |
|  | ADAR overexpression (all conditions) | EK0574 CAG-ADAR1-p150 | 30 |
| 2c | non-matching transcript | filler plasmid | 200 |
|  | matching synthetic transcript | EK0720 CMV-TO-BFPmut-3UTR(t70587-long) (FLP-IN) | 200 |
|  | original modulADAR | EK0698 SFFV-mCherry-SGc(s70587.os13)cSG-EGFP-(3xMS2) (FLP-IN) | 200 |
|  | 16nt | NSK418 SFFV-mCherry-SGc(s70587.os13.lowbBG.16bp loop)cSG-EGFP-(3xMS2) (FLP-IN) | 270 |
|  | 31nt | NSK417 SFFV-mCherry-SGc(s70587.os13.lowbBG.31bp loop)cSG-EGFP-(3xMS2) (FLP-IN) | 270 |
|  | 43nt | NSK416 SFFV-mCherry-SGc(s70587.os13.lowbBG.43bp loop)cSG-EGFP-(3xMS2) (FLP-IN) | 270 |
|  | ADAR overexpression (all conditions) | EK0574 CAG-ADAR1-p150 | 30 |
| 2d | unrelated transcript | filler plasmid | 270 |
|  | SMN exons 6, 8 | NSK162 CMV-TO-SMN2 minigene Nluc - inactive (exon 6, 8) | 270 |
|  | SMN exons 6, 7, 8 | NSK163 CMV-TO-SMN2 minigene Nluc - active (exon 6, 7, 8) | 270 |
|  | original design | NSK173 SFFV-mCherry-SGc(Smn2ex7.os13-72bp)cSG-EGFP-(3xMS2) (FLP-IN) | 200 |
|  | low background design | NSK477 SFFV-mCherry-SGc(Smn2ex7.os13-72bp.lowbBG.16bp loop)cSG-EGFP-(3xMS2) (FLP-IN) | 200 |
|  | ADAR overexpression (all conditions) | EK0574 CAG-ADAR1-p150 | 30 |
| 2f | unrelated transcript | filler plasmid | 270 |
|  | SMN exons 6, 8 | NSK162 CMV-TO-SMN2 minigene Nluc - inactive (exon 6, 8) | 270 |
|  | SMN exons 6, 7, 8 | NSK163 CMV-TO-SMN2 minigene Nluc - active (exon 6, 7, 8) | 270 |
|  | 72nt | NSK242 SFFV-mCherry-SGc(os13.NOT-SMN2.36,55,36)cSG-EGFP-(3xMS2) (FLP-IN) | 200 |
|  | 90nt | NSK241 SFFV-mCherry-SGc(os13.NOT-SMN2.45,55,45)cSG-EGFP-(3xMS2) (FLP-IN) | 200 |
|  | ADAR overexpression (all conditions) | EK0574 CAG-ADAR1-p150 | 30 |
| 2g | unrelated transcript | filler plasmid | 270 |
|  | SCN1a exons 20, 20N, 21 | NSK250 CMV-TO-SCN1A (exons 20, 20N, 21) | 270 |
|  | SCN1a exons 20, 21 | NSK249 CMV-TO-SCN1A (exons 20, 21) | 270 |
|  | 72nt | NSK253 SFFV-mCherry-SGc(os13.NOT-SCN1A_20N.36,64,36)cSG-EGFP-(3xMS2) (FLP-IN) | 200 |
|  | 90nt | NSK252 SFFV-mCherry-SGc(os13.NOT-SCN1A_20N.45,64,45)cSG-EGFP-(3xMS2) (FLP-IN) | 200 |
|  | 102nt | NSK251 SFFV-mCherry-SGc(os13.NOT-SCN1A_20N.51,64,51)cSG-EGFP-(3xMS2) (FLP-IN) | 200 |
|  | ADAR overexpression (all conditions) | EK0574 CAG-ADAR1-p150 | 30 |
| 2h | unrelated transcript | filler plasmid | 270 |
|  | SCN1a exons 20, 20N, 21 | NSK250 CMV-TO-SCN1A (exons 20, 20N, 21) | 270 |
|  | SCN1a exons 20, 21 | NSK249 CMV-TO-SCN1A (exons 20, 21) | 270 |
|  | original design | NSK252 SFFV-mCherry-SGc(os13.NOT-SCN1A_20N.45,64,45)cSG-EGFP-(3xMS2) (FLP-IN) | 200 |
|  | low background design | NSK465 SFFV-mCherry-SGc(os13.NOT-SCN1A_20N.45,64,45.lowBG.12bp loop)cSG-EGFP-(3xMS2) (FLP-IN) | 200 |
|  | ADAR overexpression (all conditions) | EK0574 CAG-ADAR1-p150 | 30 |

**Supplementary Table 2.** Details of transfections for supplementary figures.

| Figure | Description | Plasmid | Amount (ng) |
| --- | --- | --- | --- |
| 2a | non-matching transcript | filler plasmid | 270 |
|  | EBER1 | NSK176 U6TO EBER1 | 270 |
|  | modulADAR 1 | NSK178 SFFV-mCherry-SGc(EBER1.os13.1)cSG-EGFP-(3xMS2) (FLP-IN) | 200 |
|  | modulADAR 2 | NSK179 SFFV-mCherry-SGc(EBER1.os13.2)cSG-EGFP-(3xMS2) (FLP-IN) | 200 |
|  | modulADAR 3 | NSK180 SFFV-mCherry-SGc(EBER1.os13.3)cSG-EGFP-(3xMS2) (FLP-IN) | 200 |
|  | modulADAR 4 | NSK181 SFFV-mCherry-SGc(EBER1.os13.4)cSG-EGFP-(3xMS2) (FLP-IN) | 200 |
|  | modulADAR 5 | NSK182 SFFV-mCherry-SGc(EBER1.os13.5)cSG-EGFP-(3xMS2) (FLP-IN) | 200 |
|  | modulADAR 6 | NSK183 SFFV-mCherry-SGc(EBER1.os13.6)cSG-EGFP-(3xMS2) (FLP-IN) | 200 |
|  | modulADAR 7 | NSK184 SFFV-mCherry-SGc(EBER1.os13.7)cSG-EGFP-(3xMS2) (FLP-IN) | 200 |
|  | modulADAR 8 | NSK185 SFFV-mCherry-SGc(EBER1.os13.8)cSG-EGFP-(3xMS2) (FLP-IN) | 200 |
|  | modulADAR 9 | NSK186 SFFV-mCherry-SGc(EBER1.os13.9)cSG-EGFP-(3xMS2) (FLP-IN) | 200 |
|  | linear 1 | NSK265 SFFV-mCherry-SGc(EBER1.linear.1)cSG-EGFP-(3xMS2) (FLP-IN) | 200 |
|  | linear 2 | NSK266 SFFV-mCherry-SGc(EBER1.linear.2)cSG-EGFP-(3xMS2) (FLP-IN) | 200 |
|  | ADAR overexpression (all conditions) | EK0574 CAG-ADAR1-p150 | 30 |
| 2b | non-matching transcript | filler plasmid | 270 |
|  | EBER2 | NSK177 U6TO EBER2 | 270 |
|  | modulADAR 1 | NSK187 SFFV-mCherry-SGc(EBER2.os13.1)cSG-EGFP-(3xMS2) (FLP-IN) | 200 |
|  | modulADAR 2 | NSK188 SFFV-mCherry-SGc(EBER2.os13.2)cSG-EGFP-(3xMS2) (FLP-IN) | 200 |
|  | modulADAR 3 | NSK189 SFFV-mCherry-SGc(EBER2.os13.3)cSG-EGFP-(3xMS2) (FLP-IN) | 200 |
|  | modulADAR 4 | NSK190 SFFV-mCherry-SGc(EBER2.os13.4)cSG-EGFP-(3xMS2) (FLP-IN) | 200 |
|  | modulADAR 5 | NSK191 SFFV-mCherry-SGc(EBER2.os13.5)cSG-EGFP-(3xMS2) (FLP-IN) | 200 |
|  | modulADAR 6 | NSK192 SFFV-mCherry-SGc(EBER2.os13.6)cSG-EGFP-(3xMS2) (FLP-IN) | 200 |
|  | modulADAR 7 | NSK193 SFFV-mCherry-SGc(EBER2.os13.7)cSG-EGFP-(3xMS2) (FLP-IN) | 200 |
|  | modulADAR 8 | NSK194 SFFV-mCherry-SGc(EBER2.os13.8)cSG-EGFP-(3xMS2) (FLP-IN) | 200 |
|  | modulADAR 9 | NSK195 SFFV-mCherry-SGc(EBER2.os13.9)cSG-EGFP-(3xMS2) (FLP-IN) | 200 |
|  | modulADAR 10 | NSK196 SFFV-mCherry-SGc(EBER2.os13.10)cSG-EGFP-(3xMS2) (FLP-IN) | 200 |
|  | linear 1 | NSK267 SFFV-mCherry-SGc(EBER2.linear.1)cSG-EGFP-(3xMS2) (FLP-IN) | 200 |
|  | ADAR overexpression (all conditions) | EK0574 CAG-ADAR1-p150 | 30 |
| 3 | sensor | EK0438 SFFV-mCherry-SGc(s70587)cSG-EGFP-(3xMS2) (FLP-IN) | 200 |
|  | AAA | EK0580 CMV-TO-BFPmut-3UTR(t70587-AAA) (FLP-IN) | 250 |
|  | AAT | EK0581 CMV-TO-BFPmut-3UTR(t70587-AAT) (FLP-IN) | 250 |
|  | AAG | EK0582 CMV-TO-BFPmut-3UTR(t70587-AAG) (FLP-IN) | 250 |
|  | AAC | EK0583 CMV-TO-BFPmut-3UTR(t70587-AAC) (FLP-IN) | 250 |
|  | ATA | EK0584 CMV-TO-BFPmut-3UTR(t70587-ATA) (FLP-IN) | 250 |
|  | ATT | EK0585 CMV-TO-BFPmut-3UTR(t70587-ATT) (FLP-IN) | 250 |
|  | ATG | EK0586 CMV-TO-BFPmut-3UTR(t70587-ATG) (FLP-IN) | 250 |
|  | ATC | EK0587 CMV-TO-BFPmut-3UTR(t70587-ATC) (FLP-IN) | 250 |
|  | AGA | EK0588 CMV-TO-BFPmut-3UTR(t70587-AGA) (FLP-IN) | 250 |
|  | AGT | EK0589 CMV-TO-BFPmut-3UTR(t70587-AGT) (FLP-IN) | 250 |
|  | AGG | EK0590 CMV-TO-BFPmut-3UTR(t70587-AGG) (FLP-IN) | 250 |
|  | AGC | EK0591 CMV-TO-BFPmut-3UTR(t70587-AGC) (FLP-IN) | 250 |
|  | ACA | EK0592 CMV-TO-BFPmut-3UTR(t70587-ACA) (FLP-IN) | 250 |
|  | ACT | EK0593 CMV-TO-BFPmut-3UTR(t70587-ACT) (FLP-IN) | 250 |
|  | ACG | EK0594 CMV-TO-BFPmut-3UTR(t70587-ACG) (FLP-IN) | 250 |
|  | ACC | EK0595 CMV-TO-BFPmut-3UTR(t70587-ACC) (FLP-IN) | 250 |
|  | TAA | EK0596 CMV-TO-BFPmut-3UTR(t70587-TAA) (FLP-IN) | 250 |
|  | TAT | EK0597 CMV-TO-BFPmut-3UTR(t70587-TAT) (FLP-IN) | 250 |
|  | TAG | EK0598 CMV-TO-BFPmut-3UTR(t70587-TAG) (FLP-IN) | 250 |
|  | TAC | EK0599 CMV-TO-BFPmut-3UTR(t70587-TAC) (FLP-IN) | 250 |
|  | TTA | EK0600 CMV-TO-BFPmut-3UTR(t70587-TTA) (FLP-IN) | 250 |
|  | TTT | EK0601 CMV-TO-BFPmut-3UTR(t70587-TTT) (FLP-IN) | 250 |
|  | TTG | EK0602 CMV-TO-BFPmut-3UTR(t70587-TTG) (FLP-IN) | 250 |
|  | TTC | EK0603 CMV-TO-BFPmut-3UTR(t70587-TTC) (FLP-IN) | 250 |
|  | TGA | EK0604 CMV-TO-BFPmut-3UTR(t70587-TGA) (FLP-IN) | 250 |
|  | TGT | EK0605 CMV-TO-BFPmut-3UTR(t70587-TGT) (FLP-IN) | 250 |
|  | TGG | EK0606 CMV-TO-BFPmut-3UTR(t70587-TGG) (FLP-IN) | 250 |
|  | TGC | EK0607 CMV-TO-BFPmut-3UTR(t70587-TGC) (FLP-IN) | 250 |
|  | TCA | EK0608 CMV-TO-BFPmut-3UTR(t70587-TCA) (FLP-IN) | 250 |
|  | TCT | EK0609 CMV-TO-BFPmut-3UTR(t70587-TCT) (FLP-IN) | 250 |
|  | TCG | EK0610 CMV-TO-BFPmut-3UTR(t70587-TCG) (FLP-IN) | 250 |
|  | TCC | EK0611 CMV-TO-BFPmut-3UTR(t70587-TCC) (FLP-IN) | 250 |
|  | GAA | EK0612 CMV-TO-BFPmut-3UTR(t70587-GAA) (FLP-IN) | 250 |
|  | GAT | EK0613 CMV-TO-BFPmut-3UTR(t70587-GAT) (FLP-IN) | 250 |
|  | GAG | EK0614 CMV-TO-BFPmut-3UTR(t70587-GAG) (FLP-IN) | 250 |
|  | GAC | EK0615 CMV-TO-BFPmut-3UTR(t70587-GAC) (FLP-IN) | 250 |
|  | GTA | EK0616 CMV-TO-BFPmut-3UTR(t70587-GTA) (FLP-IN) | 250 |
|  | GTT | EK0617 CMV-TO-BFPmut-3UTR(t70587-GTT) (FLP-IN) | 250 |
|  | GTG | EK0618 CMV-TO-BFPmut-3UTR(t70587-GTG) (FLP-IN) | 250 |
|  | GTC | EK0619 CMV-TO-BFPmut-3UTR(t70587-GTC) (FLP-IN) | 250 |
|  | GGA | EK0620 CMV-TO-BFPmut-3UTR(t70587-GGA) (FLP-IN) | 250 |
|  | GGT | EK0621 CMV-TO-BFPmut-3UTR(t70587-GGT) (FLP-IN) | 250 |
|  | GGG | EK0622 CMV-TO-BFPmut-3UTR(t70587-GGG) (FLP-IN) | 250 |
|  | GGC | EK0623 CMV-TO-BFPmut-3UTR(t70587-GGC) (FLP-IN) | 250 |
|  | GCA | EK0624 CMV-TO-BFPmut-3UTR(t70587-GCA) (FLP-IN) | 250 |
|  | GCT | EK0625 CMV-TO-BFPmut-3UTR(t70587-GCT) (FLP-IN) | 250 |
|  | GCG | EK0626 CMV-TO-BFPmut-3UTR(t70587-GCG) (FLP-IN) | 250 |
|  | GCC | EK0627 CMV-TO-BFPmut-3UTR(t70587-GCC) (FLP-IN) | 250 |
|  | CAA | EK0628 CMV-TO-BFPmut-3UTR(t70587-CAA) (FLP-IN) | 250 |
|  | CAT | EK0629 CMV-TO-BFPmut-3UTR(t70587-CAT) (FLP-IN) | 250 |
|  | CAG | EK0630 CMV-TO-BFPmut-3UTR(t70587-CAG) (FLP-IN) | 250 |
|  | CAC | EK0631 CMV-TO-BFPmut-3UTR(t70587-CAC) (FLP-IN) | 250 |
|  | CTA | EK0632 CMV-TO-BFPmut-3UTR(t70587-CTA) (FLP-IN) | 250 |
|  | CTT | EK0633 CMV-TO-BFPmut-3UTR(t70587-CTT) (FLP-IN) | 250 |
|  | CTG | EK0634 CMV-TO-BFPmut-3UTR(t70587-CTG) (FLP-IN) | 250 |
|  | CTC | EK0635 CMV-TO-BFPmut-3UTR(t70587-CTC) (FLP-IN) | 250 |
|  | CGA | EK0636 CMV-TO-BFPmut-3UTR(t70587-CGA) (FLP-IN) | 250 |
|  | CGT | EK0637 CMV-TO-BFPmut-3UTR(t70587-CGT) (FLP-IN) | 250 |
|  | CGG | EK0638 CMV-TO-BFPmut-3UTR(t70587-CGG) (FLP-IN) | 250 |
|  | CGC | EK0639 CMV-TO-BFPmut-3UTR(t70587-CGC) (FLP-IN) | 250 |
|  | CCT | EK0640 CMV-TO-BFPmut-3UTR(t70587-CCT) (FLP-IN) | 250 |
|  | CCG | EK0641 CMV-TO-BFPmut-3UTR(t70587-CCG) (FLP-IN) | 250 |
|  | CCC | EK0642 CMV-TO-BFPmut-3UTR(t70587-CCC) (FLP-IN) | 250 |
|  | - | Filler plasmid | 250 |
|  | ADAR overexpression (all conditions) | pCDH 3FLAG-ADAR1-p150-HIS | 50 |
| 4 | non-matching transcript | filler plasmid | 270 |
|  | matching synthetic transcript | EK0720 CMV-TO-BFPmut-3UTR(t70587-long) (FLP-IN) | 270 |
|  | original sensor | EK0698 SFFV-mCherry-SGc(s70587.os13)cSG-EGFP-(3xMS2) (FLP-IN) | 200 |
|  | variant 1 | NSK231 SFFV-mCherry-SGc(s70587.os13.2stop.1)cSG-EGFP-(3xMS2) (FLP-IN) | 200 |
|  | variant 2 | NSK232 SFFV-mCherry-SGc(s70587.os13.2stop.2)cSG-EGFP-(3xMS2) (FLP-IN) | 200 |
|  | ADAR overexpression (all conditions) | EK0574 CAG-ADAR1-p150 | 30 |
| 5a | non-matching transcript | Filler plasmid | 200 |
|  | matching synthetic transcript | EK0720 CMV-TO-BFPmut-3UTR(t70587-long) (FLP-IN) | 200 |
|  | 4nt | NSK437 SFFV-mCherry-SGc(s70587.os13.lowbBG.4bp loop)cSG-EGFP-(3xMS2) (FLP-IN) | 270 |
|  | 7nt | NSK436 SFFV-mCherry-SGc(s70587.os13.lowbBG.7bp loop)cSG-EGFP-(3xMS2) (FLP-IN) | 270 |
|  | 10nt | NSK435 SFFV-mCherry-SGc(s70587.os13.lowbBG.10bp loop)cSG-EGFP-(3xMS2) (FLP-IN) | 270 |
|  | 13nt | NSK434 SFFV-mCherry-SGc(s70587.os13.lowbBG.13bp loop)cSG-EGFP-(3xMS2) (FLP-IN) | 270 |
|  | 16nt | NSK418 SFFV-mCherry-SGc(s70587.os13.lowbBG.16bp loop)cSG-EGFP-(3xMS2) (FLP-IN) | 270 |
|  | linear sensor | EK0438 SFFV-mCherry-SGc(s70587)cSG-EGFP-(3xMS2) (FLP-IN) | 270 |
|  | ADAR overexpression (all conditions) | EK0574 CAG-ADAR1-p150 | 30 |
| 5b | non-matching transcript | Filler plasmid | 270 |
|  | matching synthetic transcript | EK0720 CMV-TO-BFPmut-3UTR(t70587-long) (FLP-IN) | 270 |
|  | 13nt (contiguous) | NSK434 SFFV-mCherry-SGc(s70587.os13.lowbBG.13bp loop)cSG-EGFP-(3xMS2) (FLP-IN) | 200 |
|  | 16nt | NSK446 SFFV-mCherry-SGc(s70587.os13.lowbBG.split 1'-2' 16bp loop)cSG-EGFP-(3xMS2) (FLP-IN) | 200 |
|  | 34nt | NSK445 SFFV-mCherry-SGc(s70587.os13.lowbBG.split 1'-2' 34bp loop)cSG-EGFP-(3xMS2) (FLP-IN) | 200 |
|  | 52nt | NSK444 SFFV-mCherry-SGc(s70587.os13.lowbBG.split 1'-2' 52bp loop)cSG-EGFP-(3xMS2) (FLP-IN) | 200 |
|  | 70nt | NSK443 SFFV-mCherry-SGc(s70587.os13.lowbBG.split 1'-2' 70bp loop)cSG-EGFP-(3xMS2) (FLP-IN) | 200 |
|  | 88nt | NSK442 SFFV-mCherry-SGc(s70587.os13.lowbBG.split 1'-2' 88bp loop)cSG-EGFP-(3xMS2) (FLP-IN) | 200 |
|  | linear sensor | EK0438 SFFV-mCherry-SGc(s70587)cSG-EGFP-(3xMS2) (FLP-IN) | 200 |
|  | ADAR overexpression (all conditions) | EK0574 CAG-ADAR1-p150 | 30 |
| 5c | non-matching transcript | filler plasmid | 270 |
|  | matching synthetic transcript | EK0720 CMV-TO-BFPmut-3UTR(t70587-long) (FLP-IN) | 270 |
|  | 13nt (contiguous) | NSK434 SFFV-mCherry-SGc(s70587.os13.lowbBG.13bp loop)cSG-EGFP-(3xMS2) (FLP-IN) | 200 |
|  | 16nt | NSK451 SFFV-mCherry-SGc(s70587.os13.lowbBG.split 2'-1' 16bp loop)cSG-EGFP-(3xMS2) (FLP-IN) | 200 |
|  | 34nt | NSK450 SFFV-mCherry-SGc(s70587.os13.lowbBG.split 2'-1' 34bp loop)cSG-EGFP-(3xMS2) (FLP-IN) | 200 |
|  | 52nt | NSK449 SFFV-mCherry-SGc(s70587.os13.lowbBG.split 2'-1' 52bp loop)cSG-EGFP-(3xMS2) (FLP-IN) | 200 |
|  | 70nt | NSK448 SFFV-mCherry-SGc(s70587.os13.lowbBG.split 2'-1' 70bp loop)cSG-EGFP-(3xMS2) (FLP-IN) | 200 |
|  | 88nt | NSK447 SFFV-mCherry-SGc(s70587.os13.lowbBG.split 2'-1' 88bp loop)cSG-EGFP-(3xMS2) (FLP-IN) | 200 |
|  | linear sensor | EK0438 SFFV-mCherry-SGc(s70587)cSG-EGFP-(3xMS2) (FLP-IN) | 200 |
|  | ADAR overexpression (all conditions) | EK0574 CAG-ADAR1-p150 | 30 |
| 6a | unrelated transcript | filler plasmid | 270 |
|  | SMN exons 6, 8 | NSK162 CMV-TO-SMN2 minigene Nluc - inactive (exon 6, 8) | 270 |
|  | SMN exons 6, 7, 8 | NSK163 CMV-TO-SMN2 minigene Nluc - active (exon 6, 7, 8) | 270 |
|  | 7nt | NSK469 SFFV-mCherry-SGc(Smn2ex7.os13-72bp.lowBG.7bp loop)cSG-EGFP-(3xMS2) (FLP-IN) | 200 |
|  | 10nt | NSK468 SFFV-mCherry-SGc(Smn2ex7.os13-72bp.lowBG.10bp loop)cSG-EGFP-(3xMS2) (FLP-IN) | 200 |
|  | 13nt | NSK467 SFFV-mCherry-SGc(Smn2ex7.os13-72bp.lowBG.13bp loop)cSG-EGFP-(3xMS2) (FLP-IN) | 200 |
|  | ADAR overexpression (all conditions) | EK0574 CAG-ADAR1-p150 | 30 |
| 6b | unrelated transcript | filler plasmid | 270 |
|  | SMN exons 6, 8 | NSK162 CMV-TO-SMN2 minigene Nluc - inactive (exon 6, 8) | 270 |
|  | SMN exons 6, 7, 8 | NSK163 CMV-TO-SMN2 minigene Nluc - active (exon 6, 7, 8) | 270 |
|  | 13nt | NSK467 SFFV-mCherry-SGc(Smn2ex7.os13-72bp.lowBG.13bp loop)cSG-EGFP-(3xMS2) (FLP-IN) | 200 |
|  | 16nt | NSK477 SFFV-mCherry-SGc(Smn2ex7.os13-72bp.lowBG.16bp loop)cSG-EGFP-(3xMS2) (FLP-IN) | 200 |
|  | 19nt | NSK476 SFFV-mCherry-SGc(Smn2ex7.os13-72bp.lowBG.19bp loop)cSG-EGFP-(3xMS2) (FLP-IN) | 200 |
|  | 22nt | NSK475 SFFV-mCherry-SGc(Smn2ex7.os13-72bp.lowBG.22bp loop)cSG-EGFP-(3xMS2) (FLP-IN) | 200 |
|  | ADAR overexpression (all conditions) | EK0574 CAG-ADAR1-p150 | 30 |
| **7b** | unrelated transcript | filler plasmid | 270 |
|  | SCN1a exons 20, 20N, 21 | NSK250 CMV-TO-SCN1A (exons 20, 20N, 21) | 270 |
|  | SCN1a exons 20, 21 | NSK249 CMV-TO-SCN1A (exons 20, 21) | 270 |
|  | original sensor | NSK252 SFFV-mCherry-SGc(os13.NOT-SCN1A_20N.45,64,45)cSG-EGFP-(3xMS2) (FLP-IN) | 200 |
|  | 6nt | NSK466 SFFV-mCherry-SGc(os13.NOT-SCN1A_20N.45,64,45.lowBG.6bp loop)cSG-EGFP-(3xMS2) (FLP-IN) | 200 |
|  | 12nt | NSK465 SFFV-mCherry-SGc(os13.NOT-SCN1A_20N.45,64,45.lowBG.12bp loop)cSG-EGFP-(3xMS2) (FLP-IN) | 200 |
|  | 18nt | NSK464 SFFV-mCherry-SGc(os13.NOT-SCN1A_20N.45,64,45.lowBG.18bp loop)cSG-EGFP-(3xMS2) (FLP-IN) | 200 |
|  | ADAR overexpression (all conditions) | EK0574 CAG-ADAR1-p150 | 30 |
| **7c** | unrelated transcript | filler plasmid | 270 |
|  | SCN1a exons 20, 20N, 21 | NSK250 CMV-TO-SCN1A (exons 20, 20N, 21) | 270 |
|  | SCN1a exons 20, 21 | NSK249 CMV-TO-SCN1A (exons 20, 21) | 270 |
|  | 9nt | NSK482 SFFV-mCherry-SGc(os13.NOT-SCN1A_20N.45,64,45.lowBG.9bp loop)cSG-EGFP-(3xMS2) (FLP-IN) | 200 |
|  | 12nt | NSK465 SFFV-mCherry-SGc(os13.NOT-SCN1A_20N.45,64,45.lowBG.12bp loop)cSG-EGFP-(3xMS2) (FLP-IN) | 200 |
|  | 15nt | NSK481 SFFV-mCherry-SGc(os13.NOT-SCN1A_20N.45,64,45.lowBG.15bp loop)cSG-EGFP-(3xMS2) (FLP-IN) | 200 |
|  | ADAR overexpression (all conditions) | EK0574 CAG-ADAR1-p150 | 30 |

**Supplementary Table 3.** List of plasmids used in main text figures. HIghlighted plasmids are deposited on addgene. Other plasmids are available upon reasonable request.

| Plasmid name | Source | Addgene ID | Map |
| --- | --- | --- | --- |
| EK0301 SFFV-Che-SG(Bdnf.s1)SG-GFP (FLP-IN) | Prior work | 191144 | benchling link |
| EK0307 CMV-TO-BFPmut-stop-3UTR(Bdnf)-v3 (FLP-IN) | Prior work | 191145 | benchling link |
| EK0438 SFFV-mCherry-SGc(s70587)cSG-EGFP-(3xMS2) (FLP-IN) | Prior work | 191163 | benchling link |
| EK0574 CAG-ADAR1-p150 | This work | TBD | benchling link |
| EK0656 SFFV-mCherry-SGc(70587.os4)cSG-EGFP-(3xMS2) | This work | TBD | benchling link |
| EK0698 SFFV-mCherry-SGc(s70587.os13)cSG-EGFP-(3xMS2) (FLP-IN) | This work | TBD | benchling link |
| EK0706 SFFV-mCherry-SGc(Bdnf.os4)cSG-EGFP-(3xMS2) (FLP-IN) | This work | NA | benchling link |
| EK0716 SFFV-mCherry-SGc(s70587.os18)cSG-EGFP-(3xMS2) (FLP-IN) | This work | TBD | benchling link |
| EK0717 SFFV-mCherry-SGc(s70587.os19)cSG-EGFP-(3xMS2) (FLP-IN) | This work | TBD | benchling link |
| EK0718 SFFV-mCherry-SGc(s70587.os20)cSG-EGFP-(3xMS2) (FLP-IN) | This work | TBD | benchling link |
| EK0720 CMV-TO-BFPmut-3UTR(t70587-long) (FLP-IN) | This work | TBD | benchling link |
| NSK157 SFFV-mCherry-SGc(s70587.os-360bp)cSG-EGFP-(3xMS2) (FLP-IN) | This work | NA | benchling link |
| NSK158 SFFV-mCherry-SGc(s70587.os-180bp)cSG-EGFP-(3xMS2) (FLP-IN) | This work | NA | benchling link |
| NSK159 SFFV-mCherry-SGc(s70587.os-102bp)cSG-EGFP-(3xMS2) (FLP-IN) | This work | NA | benchling link |
| NSK160 SFFV-mCherry-SGc(s70587.os-60bp)cSG-EGFP-(3xMS2) (FLP-IN) | This work | NA | benchling link |
| NSK162 CMV-TO-SMN2 Nluc - inactive (exon 6, 8) | This work | TBD | benchling link |
| NSK163 CMV-TO-SMN2 Nluc - active (exon 6, 7, 8) | This work | TBD | benchling link |
| NSK165 SFFV-mCherry-SGc(Bdnf.os13)cSG-EGFP-(3xMS2) (FLP-IN) | This work | NA | benchling link |
| NSK166 SFFV-mCherry-SGc(Bdnf.os18)cSG-EGFP-(3xMS2) (FLP-IN) | This work | NA | benchling link |
| NSK167 SFFV-mCherry-SGc(Bdnf.os20)cSG-EGFP-(3xMS2) (FLP-IN) | This work | NA | benchling link |
| NSK171 SFFV-mCherry-SGc(Smn2ex7.os13-90bp)cSG-EGFP-(3xMS2) (FLP-IN) | This work | NA | benchling link |
| NSK172 SFFV-mCherry-SGc(Smn2ex7.os13-102bp)cSG-EGFP-(3xMS2) (FLP-IN) | This work | NA | benchling link |
| NSK173 SFFV-mCherry-SGc(Smn2ex7.os13-72bp)cSG-EGFP-(3xMS2) (FLP-IN) | This work | TBD | benchling link |
| NSK176 U6TO EBER1 | This work | TBD | benchling link |
| NSK177 U6TO EBER2 | This work | TBD | benchling link |
| NSK185 SFFV-mCherry-SGc(EBER1.os13.8)cSG-EGFP-(3xMS2) (FLP-IN) | This work | TBD | benchling link |
| NSK195 SFFV-mCherry-SGc(EBER2.os13.9)cSG-EGFP-(3xMS2) (FLP-IN) | This work | TBD | benchling link |
| NSK204 SFFV-mCherry-SGc(SNM2.scram1)cSG-EGFP-(3xMS2) (FLP-IN) | This work | NA | benchling link |
| NSK205 SFFV-mCherry-SGc(SNM2.scram2)cSG-EGFP-(3xMS2) (FLP-IN) | This work | NA | benchling link |
| NSK206 SFFV-mCherry-SGc(SNM2.scram3)cSG-EGFP-(3xMS2) (FLP-IN) | This work | NA | benchling link |
| NSK207 SFFV-mCherry-SGc(SNM2.scram4)cSG-EGFP-(3xMS2) (FLP-IN) | This work | NA | benchling link |
| NSK241 SFFV-mCherry-SGc(os13.NOT-SMN2.45,55,45)cSG-EGFP-(3xMS2) (FLP-IN) | This work | TBD | benchling link |
| NSK242 SFFV-mCherry-SGc(os13.NOT-SMN2.36,55,36)cSG-EGFP-(3xMS2) (FLP-IN) | This work | TBD | benchling link |
| NSK249 CMV-TO-SCN1A (exons 20, 21) | This work | TBD | benchling link |
| NSK250 CMV-TO-SCN1A (exons 20, 20N, 21) | This work | TBD | benchling link |
| NSK251 SFFV-mCherry-SGc(os13.NOT-SCN1A_20N.51,64,51)cSG-EGFP-(3xMS2) (FLP-IN) | This work | NA | benchling link |
| NSK252 SFFV-mCherry-SGc(os13.NOT-SCN1A_20N.45,64,45)cSG-EGFP-(3xMS2) (FLP-IN) | This work | TBD | benchling link |
| NSK253 SFFV-mCherry-SGc(os13.NOT-SCN1A_20N.36,64,36)cSG-EGFP-(3xMS2) (FLP-IN) | This work | NA | benchling link |
| NSK266 SFFV-mCherry-SGc(EBER1.linear.2)cSG-EGFP-(3xMS2) (FLP-IN) | This work | TBD | benchling link |
| NSK267 SFFV-mCherry-SGc(EBER2.linear.1)cSG-EGFP-(3xMS2) (FLP-IN) | This work | TBD | benchling link |
| NSK416 SFFV-mCherry-SGc(s70587.os13.lowbBG.43bp loop)cSG-EGFP-(3xMS2) (FLP-IN) | This work | NA | benchling link |
| NSK417 SFFV-mCherry-SGc(s70587.os13.lowbBG.31bp loop)cSG-EGFP-(3xMS2) (FLP-IN) | This work | NA | benchling link |
| NSK418 SFFV-mCherry-SGc(s70587.os13.lowbBG.16bp loop)cSG-EGFP-(3xMS2) (FLP-IN) | This work | TBD | benchling link |
| NSK465 SFFV-mCherry-SGc(os13.NOT-SCN1A_20N.45,64,45.lowBG.12bp loop)cSG-EGFP-(3xMS2) (FLP-IN) | This work | TBD | benchling link |
| NSK477 SFFV-mCherry-SGc(Smn2ex7.os13-72bp.lowBG.16bp loop)cSG-EGFP-(3xMS2) (FLP-IN) | This work | TBD | benchling link |

**Supplementary Table 4.** List of plasmids used in only in SI figures. Plasmids are available upon reasonable request.

| Plasmid name | Source | Map |
| --- | --- | --- |
| EK0580 CMV-TO-BFPmut-3UTR(t70587-AAA) (FLP-IN) | This work | benchling link |
| EK0581 CMV-TO-BFPmut-3UTR(t70587-AAT) (FLP-IN) | This work | benchling link |
| EK0582 CMV-TO-BFPmut-3UTR(t70587-AAG) (FLP-IN) | This work | benchling link |
| EK0583 CMV-TO-BFPmut-3UTR(t70587-AAC) (FLP-IN) | This work | benchling link |
| EK0584 CMV-TO-BFPmut-3UTR(t70587-ATA) (FLP-IN) | This work | benchling link |
| EK0585 CMV-TO-BFPmut-3UTR(t70587-ATT) (FLP-IN) | This work | benchling link |
| EK0586 CMV-TO-BFPmut-3UTR(t70587-ATG) (FLP-IN) | This work | benchling link |
| EK0587 CMV-TO-BFPmut-3UTR(t70587-ATC) (FLP-IN) | This work | benchling link |
| EK0588 CMV-TO-BFPmut-3UTR(t70587-AGA) (FLP-IN) | This work | benchling link |
| EK0589 CMV-TO-BFPmut-3UTR(t70587-AGT) (FLP-IN) | This work | benchling link |
| EK0590 CMV-TO-BFPmut-3UTR(t70587-AGG) (FLP-IN) | This work | benchling link |
| EK0591 CMV-TO-BFPmut-3UTR(t70587-AGC) (FLP-IN) | This work | benchling link |
| EK0592 CMV-TO-BFPmut-3UTR(t70587-ACA) (FLP-IN) | This work | benchling link |
| EK0593 CMV-TO-BFPmut-3UTR(t70587-ACT) (FLP-IN) | This work | benchling link |
| EK0594 CMV-TO-BFPmut-3UTR(t70587-ACG) (FLP-IN) | This work | benchling link |
| EK0595 CMV-TO-BFPmut-3UTR(t70587-ACC) (FLP-IN) | This work | benchling link |
| EK0596 CMV-TO-BFPmut-3UTR(t70587-TAA) (FLP-IN) | This work | benchling link |
| EK0597 CMV-TO-BFPmut-3UTR(t70587-TAT) (FLP-IN) | This work | benchling link |
| EK0598 CMV-TO-BFPmut-3UTR(t70587-TAG) (FLP-IN) | This work | benchling link |
| EK0599 CMV-TO-BFPmut-3UTR(t70587-TAC) (FLP-IN) | This work | benchling link |
| EK0600 CMV-TO-BFPmut-3UTR(t70587-TTA) (FLP-IN) | This work | benchling link |
| EK0601 CMV-TO-BFPmut-3UTR(t70587-TTT) (FLP-IN) | This work | benchling link |
| EK0602 CMV-TO-BFPmut-3UTR(t70587-TTG) (FLP-IN) | This work | benchling link |
| EK0603 CMV-TO-BFPmut-3UTR(t70587-TTC) (FLP-IN) | This work | benchling link |
| EK0604 CMV-TO-BFPmut-3UTR(t70587-TGA) (FLP-IN) | This work | benchling link |
| EK0605 CMV-TO-BFPmut-3UTR(t70587-TGT) (FLP-IN) | This work | benchling link |
| EK0606 CMV-TO-BFPmut-3UTR(t70587-TGG) (FLP-IN) | This work | benchling link |
| EK0607 CMV-TO-BFPmut-3UTR(t70587-TGC) (FLP-IN) | This work | benchling link |
| EK0608 CMV-TO-BFPmut-3UTR(t70587-TCA) (FLP-IN) | This work | benchling link |
| EK0609 CMV-TO-BFPmut-3UTR(t70587-TCT) (FLP-IN) | This work | benchling link |
| EK0610 CMV-TO-BFPmut-3UTR(t70587-TCG) (FLP-IN) | This work | benchling link |
| EK0611 CMV-TO-BFPmut-3UTR(t70587-TCC) (FLP-IN) | This work | benchling link |
| EK0612 CMV-TO-BFPmut-3UTR(t70587-GAA) (FLP-IN) | This work | benchling link |
| EK0613 CMV-TO-BFPmut-3UTR(t70587-GAT) (FLP-IN) | This work | benchling link |
| EK0614 CMV-TO-BFPmut-3UTR(t70587-GAG) (FLP-IN) | This work | benchling link |
| EK0615 CMV-TO-BFPmut-3UTR(t70587-GAC) (FLP-IN) | This work | benchling link |
| EK0616 CMV-TO-BFPmut-3UTR(t70587-GTA) (FLP-IN) | This work | benchling link |
| EK0617 CMV-TO-BFPmut-3UTR(t70587-GTT) (FLP-IN) | This work | benchling link |
| EK0618 CMV-TO-BFPmut-3UTR(t70587-GTG) (FLP-IN) | This work | benchling link |
| EK0619 CMV-TO-BFPmut-3UTR(t70587-GTC) (FLP-IN) | This work | benchling link |
| EK0620 CMV-TO-BFPmut-3UTR(t70587-GGA) (FLP-IN) | This work | benchling link |
| EK0621 CMV-TO-BFPmut-3UTR(t70587-GGT) (FLP-IN) | This work | benchling link |
| EK0622 CMV-TO-BFPmut-3UTR(t70587-GGG) (FLP-IN) | This work | benchling link |
| EK0623 CMV-TO-BFPmut-3UTR(t70587-GGC) (FLP-IN) | This work | benchling link |
| EK0624 CMV-TO-BFPmut-3UTR(t70587-GCA) (FLP-IN) | This work | benchling link |
| EK0625 CMV-TO-BFPmut-3UTR(t70587-GCT) (FLP-IN) | This work | benchling link |
| EK0626 CMV-TO-BFPmut-3UTR(t70587-GCG) (FLP-IN) | This work | benchling link |
| EK0627 CMV-TO-BFPmut-3UTR(t70587-GCC) (FLP-IN) | This work | benchling link |
| EK0628 CMV-TO-BFPmut-3UTR(t70587-CAA) (FLP-IN) | This work | benchling link |
| EK0629 CMV-TO-BFPmut-3UTR(t70587-CAT) (FLP-IN) | This work | benchling link |
| EK0630 CMV-TO-BFPmut-3UTR(t70587-CAG) (FLP-IN) | This work | benchling link |
| EK0631 CMV-TO-BFPmut-3UTR(t70587-CAC) (FLP-IN) | This work | benchling link |
| EK0632 CMV-TO-BFPmut-3UTR(t70587-CTA) (FLP-IN) | This work | benchling link |
| EK0633 CMV-TO-BFPmut-3UTR(t70587-CTT) (FLP-IN) | This work | benchling link |
| EK0634 CMV-TO-BFPmut-3UTR(t70587-CTG) (FLP-IN) | This work | benchling link |
| EK0635 CMV-TO-BFPmut-3UTR(t70587-CTC) (FLP-IN) | This work | benchling link |
| EK0636 CMV-TO-BFPmut-3UTR(t70587-CGA) (FLP-IN) | This work | benchling link |
| EK0637 CMV-TO-BFPmut-3UTR(t70587-CGT) (FLP-IN) | This work | benchling link |
| EK0638 CMV-TO-BFPmut-3UTR(t70587-CGG) (FLP-IN) | This work | benchling link |
| EK0639 CMV-TO-BFPmut-3UTR(t70587-CGC) (FLP-IN) | This work | benchling link |
| EK0640 CMV-TO-BFPmut-3UTR(t70587-CCT) (FLP-IN) | This work | benchling link |
| EK0641 CMV-TO-BFPmut-3UTR(t70587-CCG) (FLP-IN) | This work | benchling link |
| EK0642 CMV-TO-BFPmut-3UTR(t70587-CCC) (FLP-IN) | This work | benchling link |
| NSK178 SFFV-mCherry-SGc(EBER1.os13.1)cSG-EGFP-(3xMS2) (FLP-IN) | This work | benchling link |
| NSK179 SFFV-mCherry-SGc(EBER1.os13.2)cSG-EGFP-(3xMS2) (FLP-IN) | This work | benchling link |
| NSK180 SFFV-mCherry-SGc(EBER1.os13.3)cSG-EGFP-(3xMS2) (FLP-IN) | This work | benchling link |
| NSK181 SFFV-mCherry-SGc(EBER1.os13.4)cSG-EGFP-(3xMS2) (FLP-IN) | This work | benchling link |
| NSK182 SFFV-mCherry-SGc(EBER1.os13.5)cSG-EGFP-(3xMS2) (FLP-IN) | This work | benchling link |
| NSK183 SFFV-mCherry-SGc(EBER1.os13.6)cSG-EGFP-(3xMS2) (FLP-IN) | This work | benchling link |
| NSK184 SFFV-mCherry-SGc(EBER1.os13.7)cSG-EGFP-(3xMS2) (FLP-IN) | This work | benchling link |
| NSK186 SFFV-mCherry-SGc(EBER1.os13.9)cSG-EGFP-(3xMS2) (FLP-IN) | This work | benchling link |
| NSK187 SFFV-mCherry-SGc(EBER2.os13.1)cSG-EGFP-(3xMS2) (FLP-IN) | This work | benchling link |
| NSK188 SFFV-mCherry-SGc(EBER2.os13.2)cSG-EGFP-(3xMS2) (FLP-IN) | This work | benchling link |
| NSK189 SFFV-mCherry-SGc(EBER2.os13.3)cSG-EGFP-(3xMS2) (FLP-IN) | This work | benchling link |
| NSK190 SFFV-mCherry-SGc(EBER2.os13.4)cSG-EGFP-(3xMS2) (FLP-IN) | This work | benchling link |
| NSK191 SFFV-mCherry-SGc(EBER2.os13.5)cSG-EGFP-(3xMS2) (FLP-IN) | This work | benchling link |
| NSK192 SFFV-mCherry-SGc(EBER2.os13.6)cSG-EGFP-(3xMS2) (FLP-IN) | This work | benchling link |
| NSK193 SFFV-mCherry-SGc(EBER2.os13.7)cSG-EGFP-(3xMS2) (FLP-IN) | This work | benchling link |
| NSK194 SFFV-mCherry-SGc(EBER2.os13.8)cSG-EGFP-(3xMS2) (FLP-IN) | This work | benchling link |
| NSK196 SFFV-mCherry-SGc(EBER2.os13.10)cSG-EGFP-(3xMS2) (FLP-IN) | This work | benchling link |
| NSK231 SFFV-mCherry-SGc(s70587.os13.2stop.1)cSG-EGFP-(3xMS2) (FLP-IN) | This work | benchling link |
| NSK232 SFFV-mCherry-SGc(s70587.os13.2stop.2)cSG-EGFP-(3xMS2) (FLP-IN) | This work | benchling link |
| NSK265 SFFV-mCherry-SGc(EBER1.linear.1)cSG-EGFP-(3xMS2) (FLP-IN) | This work | benchling link |
| NSK434 SFFV-mCherry-SGc(s70587.os13.lowbBG.13bp loop)cSG-EGFP-(3xMS2) (FLP-IN) | This work | benchling link |
| NSK435 SFFV-mCherry-SGc(s70587.os13.lowbBG.10bp loop)cSG-EGFP-(3xMS2) (FLP-IN) | This work | benchling link |
| NSK436 SFFV-mCherry-SGc(s70587.os13.lowbBG.7bp loop)cSG-EGFP-(3xMS2) (FLP-IN) | This work | benchling link |
| NSK437 SFFV-mCherry-SGc(s70587.os13.lowbBG.4bp loop)cSG-EGFP-(3xMS2) (FLP-IN) | This work | benchling link |
| NSK442 SFFV-mCherry-SGc(s70587.os13.lowbBG.split 1'-2' 88bp loop)cSG-EGFP-(3xMS2) (FLP-IN) | This work | benchling link |
| NSK443 SFFV-mCherry-SGc(s70587.os13.lowbBG.split 1'-2' 70bp loop)cSG-EGFP-(3xMS2) (FLP-IN) | This work | benchling link |
| NSK444 SFFV-mCherry-SGc(s70587.os13.lowbBG.split 1'-2' 52bp loop)cSG-EGFP-(3xMS2) (FLP-IN) | This work | benchling link |
| NSK445 SFFV-mCherry-SGc(s70587.os13.lowbBG.split 1'-2' 34bp loop)cSG-EGFP-(3xMS2) (FLP-IN) | This work | benchling link |
| NSK446 SFFV-mCherry-SGc(s70587.os13.lowbBG.split 1'-2' 16bp loop)cSG-EGFP-(3xMS2) (FLP-IN) | This work | benchling link |
| NSK447 SFFV-mCherry-SGc(s70587.os13.lowbBG.split 2'-1' 88bp loop)cSG-EGFP-(3xMS2) (FLP-IN) | This work | benchling link |
| NSK448 SFFV-mCherry-SGc(s70587.os13.lowbBG.split 2'-1' 70bp loop)cSG-EGFP-(3xMS2) (FLP-IN) | This work | benchling link |
| NSK449 SFFV-mCherry-SGc(s70587.os13.lowbBG.split 2'-1' 52bp loop)cSG-EGFP-(3xMS2) (FLP-IN) | This work | benchling link |
| NSK450 SFFV-mCherry-SGc(s70587.os13.lowbBG.split 2'-1' 34bp loop)cSG-EGFP-(3xMS2) (FLP-IN) | This work | benchling link |
| NSK451 SFFV-mCherry-SGc(s70587.os13.lowbBG.split 2'-1' 16bp loop)cSG-EGFP-(3xMS2) (FLP-IN) | This work | benchling link |
| NSK464 SFFV-mCherry-SGc(os13.NOT-SCN1A_20N.45,64,45.lowBG.18bp loop)cSG-EGFP-(3xMS2) (FLP-IN) | This work | benchling link |
| NSK466 SFFV-mCherry-SGc(os13.NOT-SCN1A_20N.45,64,45.lowBG.6bp loop)cSG-EGFP-(3xMS2) (FLP-IN) | This work | benchling link |
| NSK467 SFFV-mCherry-SGc(Smn2ex7.os13-72bp.lowBG.13bp loop)cSG-EGFP-(3xMS2) (FLP-IN) | This work | benchling link |
| NSK468 SFFV-mCherry-SGc(Smn2ex7.os13-72bp.lowBG.10bp loop)cSG-EGFP-(3xMS2) (FLP-IN) | This work | benchling link |
| NSK469 SFFV-mCherry-SGc(Smn2ex7.os13-72bp.lowBG.7bp loop)cSG-EGFP-(3xMS2) (FLP-IN) | This work | benchling link |
| NSK475 SFFV-mCherry-SGc(Smn2ex7.os13-72bp.lowBG.22bp loop)cSG-EGFP-(3xMS2) (FLP-IN) | This work | benchling link |
| NSK476 SFFV-mCherry-SGc(Smn2ex7.os13-72bp.lowBG.19bp loop)cSG-EGFP-(3xMS2) (FLP-IN) | This work | benchling link |
| NSK481 SFFV-mCherry-SGc(os13.NOT-SCN1A_20N.45,64,45.lowBG.15bp loop)cSG-EGFP-(3xMS2) (FLP-IN) | This work | benchling link |
| NSK482 SFFV-mCherry-SGc(os13.NOT-SCN1A_20N.45,64,45.lowBG.9bp loop)cSG-EGFP-(3xMS2) (FLP-IN) | This work | benchling link |
| pCDH 3FLAG-ADAR1-p150-HIS | Gift of J.B. Li | benchling link |

